## Supplementary material for "NanoPlex: a universal strategy for fluorescence microscopy multiplexing using nanobodies with erasable signals": Supp. Fig., Supp Table

### Synthesis and Characterization of Compounds

Supplementary Figure 1 shows an overview of the synthetic approach towards the LRT as described in the methods sections. Detailed synthetic protocols and data for compound characterization are given at the end of the Supplementary Information.

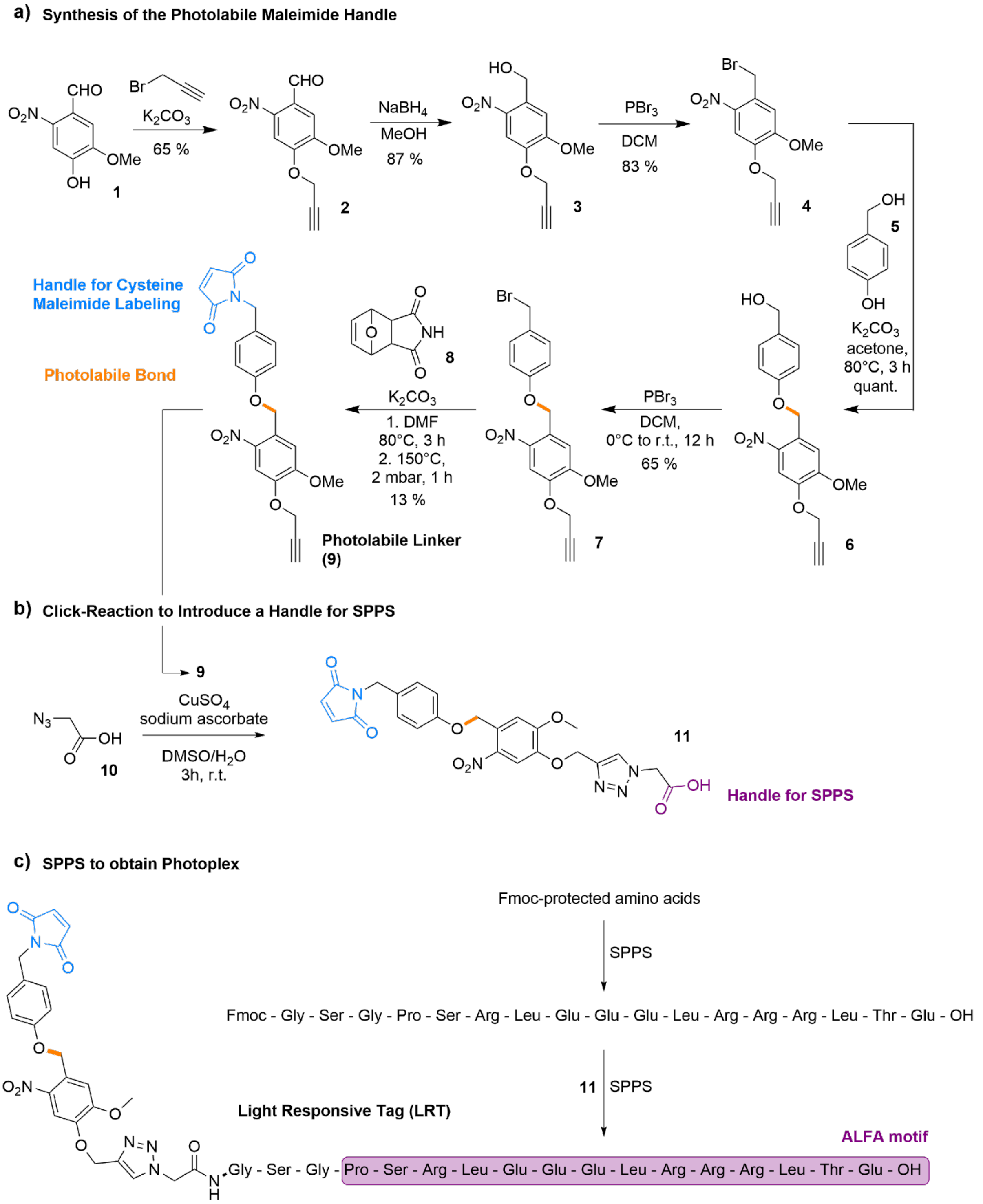


**Supplementary Figure 1.** Synthetic pathway to LRT. a: Synthesis of compound photolabile linker **9** from nitroaldehyde 1. b: click reaction to introduce a peptide handle for SPPS. c: SPPS towards LRT. More bellow in section: "**Detailed Synthetic Procedures and Compound Characterization**"

**Photophysical and photochemical characterization of LRT.**

The photophysical properties of the LRT were approximated by using the **9**, which is chemically identical to the LRT around the chromophore. The molar attenuation coefficient was determined at the lowest electronic transition around 340 nm to be 5860 L*cm-1*mol-1 (Supp. Fig. 2)

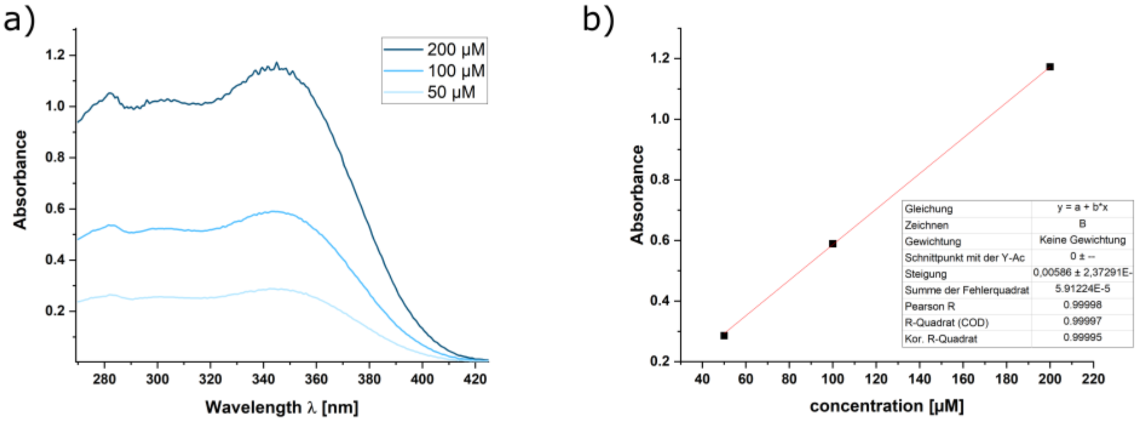


**Supplementary Figure 2**. a) Absorption spectra of **9** in DMSO at different concentrations. b) Plot of the absorption against the concentration at a specific wavelength (λ340nm) for **9** in DMSO.

To get a more detailed insight into the photochemical properties of the photolabile core, we prepared a solution of **9** (50µM, 2 mL, DMSO) and studied the photocleavage efficiency of the compound at 20 °C. We found that irradiation with a 365 nm LED (*ca.* 60mW at the outer cuvette wall) leads to the efficient disappearance of the starting material (Supp. Fig. 3a, b) and formation of the photoproduct (Supp. Fig. 3a, b). Moreover, by reducing the power of the LED, we could see that both the disappearance of the starting material and the formation of the photoproduct proceeded in a dose-dependent manner.

We also approximated the quantum yield of LRT using **9** at 100% power of the 365nm LED following a protocol that we previously adapted (Supp. Fig3a, c)[1]. Specifically, we used the initial disappearance of **9**, which was fitted (Supp. Fig. 3c) and corrected for the absorbance in the low absorption regime (Supp. Fig. 3c, d).

Next, we performed the same experiment at different wavelengths. Having set all LEDs to full power, we irradiated fresh samples of **9** (50µM, 2mL, DMSO) at 365, 405, 445, 505, and 625nm (Supp. Fig. 3e, f). As expected, photochemical bond cleavage could be induced with both 365 and 405nm; however, at 405nm the observed conversion was slower due to **9** being less absorbing at this wavelength. Beyond the absorption spectrum of 9, i.e., at a wavelength of 445nm and higher, no photoconversion was observed even at extended irradiation times (Supp. Fig. 3f).

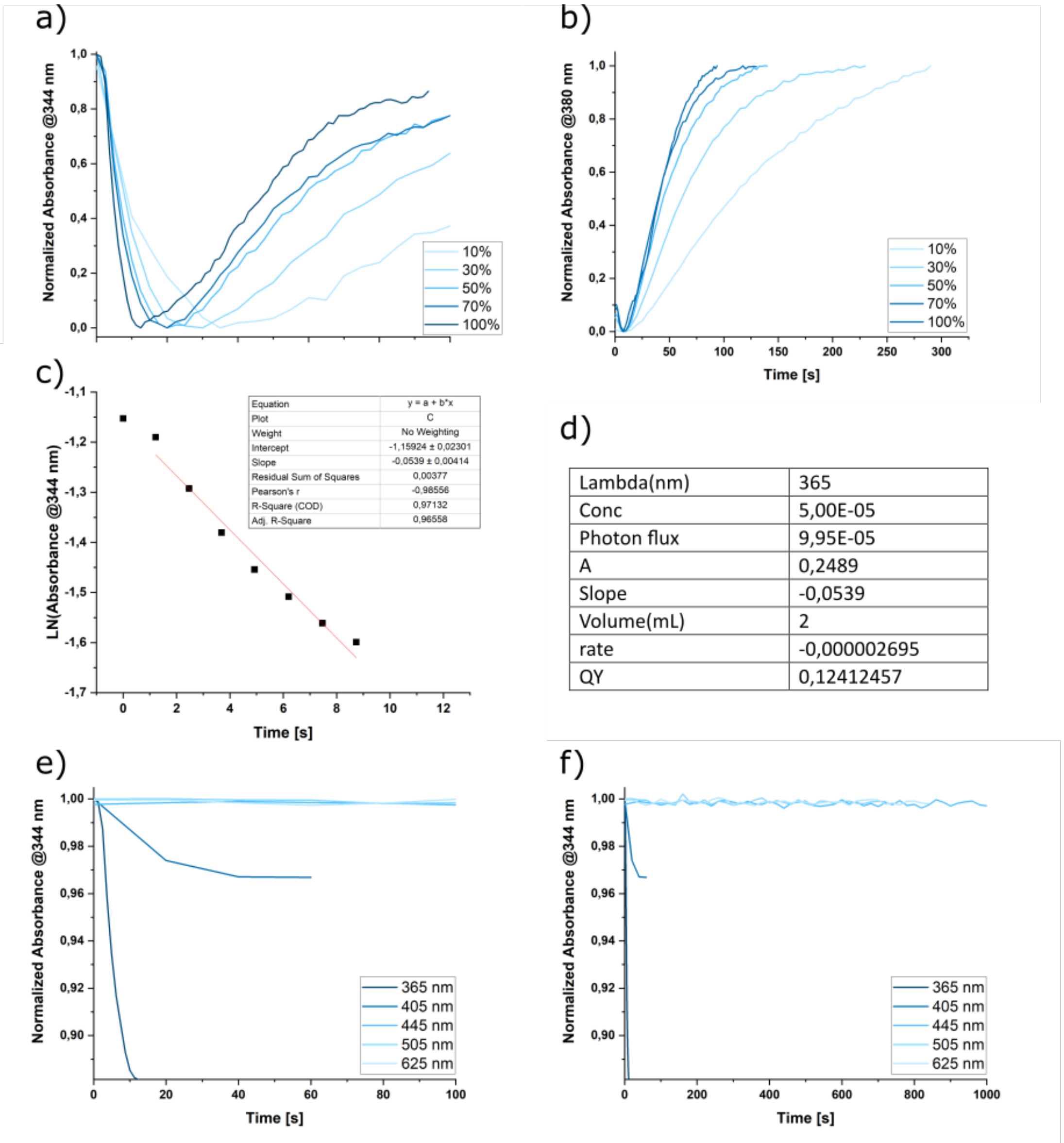


| | |
|---|---|
| Lambda(nm) | 365 |
| Conc | 5,00E-05 |
| Photon flux | 9,95E-05 |
| A | 0,2489 |
| Slope | -0,0539 |
| Volume(mL) | 2 |
| rate | -0,000002695 |
| QY | 0,12412457 |

**Supplementary Figure 3.** a) The normalized absorbance of a solution of **9** (50µM, 2mL, DMSO) at 344 nm at different irradiation intensities is shown. The decrease of the starting material is over time, overlapping with the formation of the photoproduct absorbing at the same wavelength range. b) The normalized absorbance of a solution of **9** (50µM, 2 mL, DMSO) at 380nm at different irradiation intensities is shown. The increase of the product absorption over time is shown (the spectrum is initially decreasing due to the disappearance of starting material visible at the same wavelength). c) The linearization of the absorption trace of a solution of **9** (50µM, 2mL, DMSO) at 344nm is shown. d) Parameters used to calculate the quantum yield of photochemical cleavage of **9**. e) The normalized absorbance of a solution of **9** (50µM, 2mL, DMSO) at 344nm at different wavelengths of irradiation is shown. f) The normalized absorbance of a solution of **9** (50µM, 2 mL, DMSO) at 344nm at different wavelengths of irradiation over the first 100 seconds is shown.

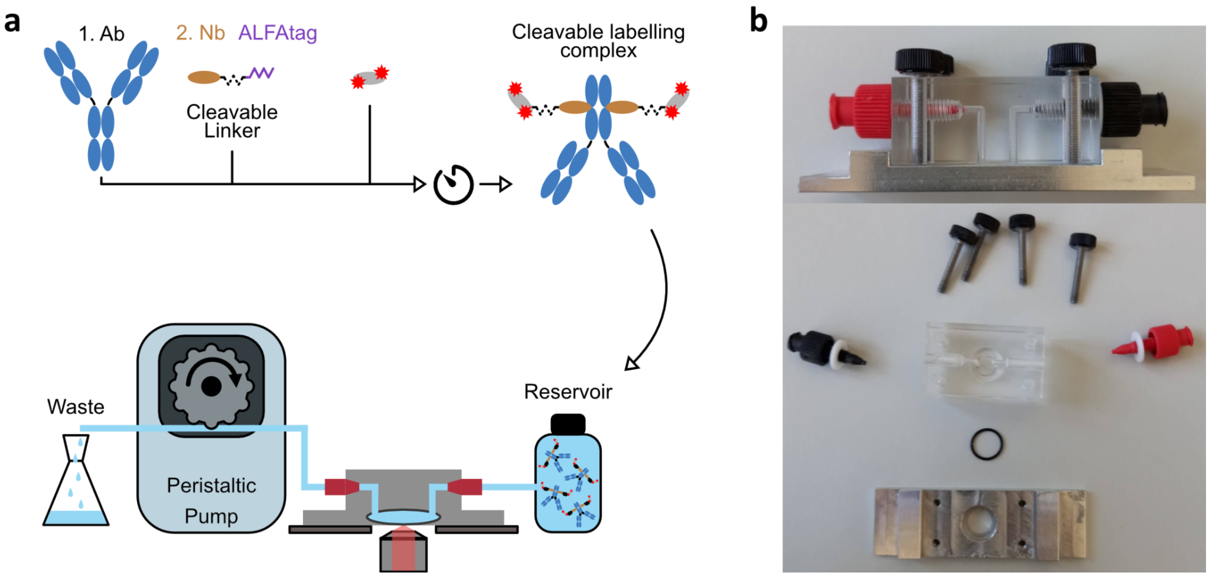


**Supplementary Figure 4: NanoPlex. a,** One-step IF and scheme of the fluidic system to perform staining and washings. **b,** Photograph of the custom-made flow chamber to use cells grown on coverslips.

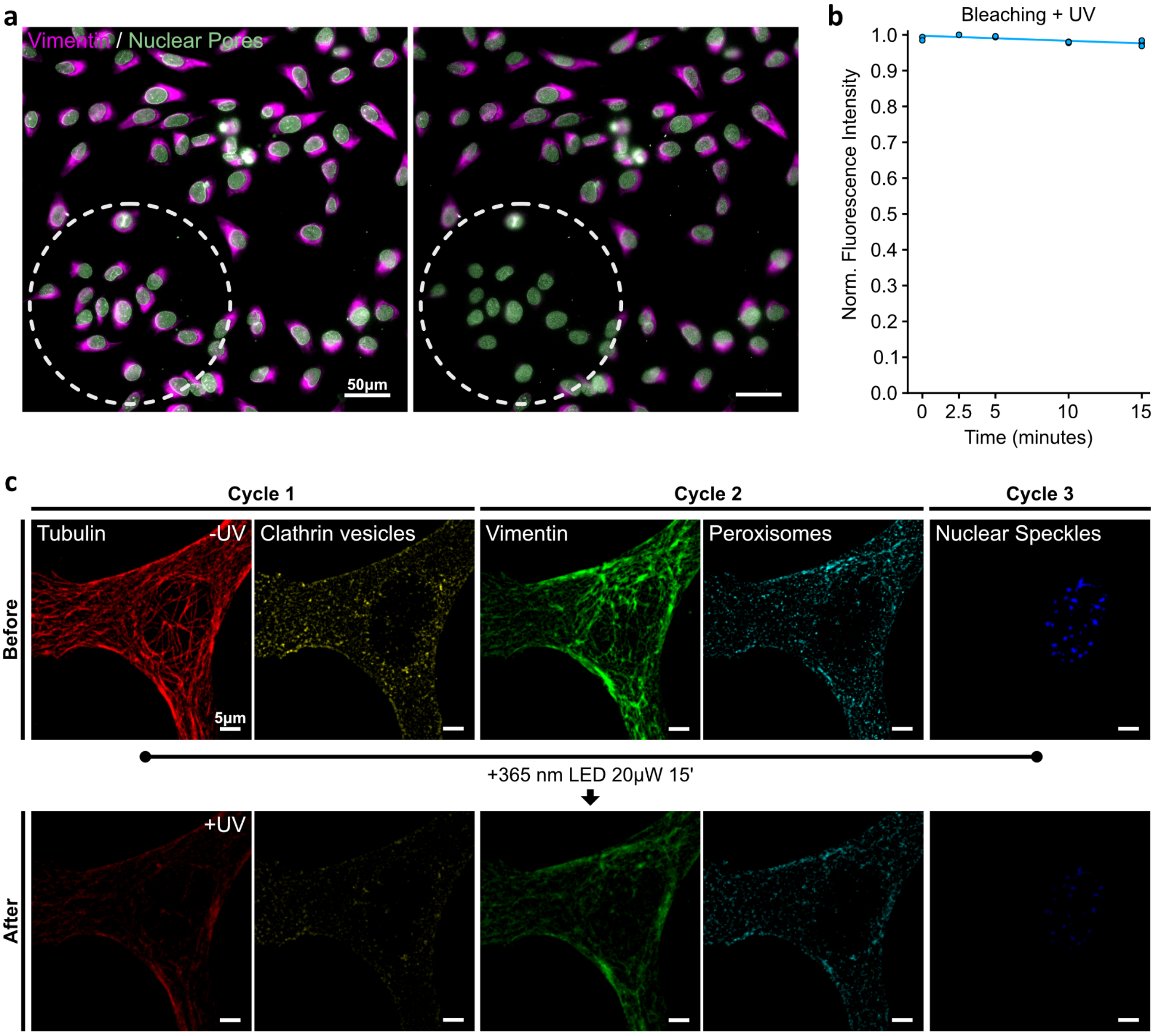


**Supplementary Figure 5. OptoPlex cleaving assessment. a,** Larger field of view, including the region (dotted circle) where the sample was illuminated with 365nm LED light for 15 minutes. **b,** Normalized Intensity of images taken while continuously illuminating the sample with 20µW of UV light (365nm) for 15 minutes. Three independent samples were stained using 2.Nbs directly labeled with Atto643 (not cleavable). **c**, full field of view of confocal images before and after illumination with 365nm light for 15 minutes. Images were acquired with equal settings and displayed with equally scaled gray levels for direct comparison.

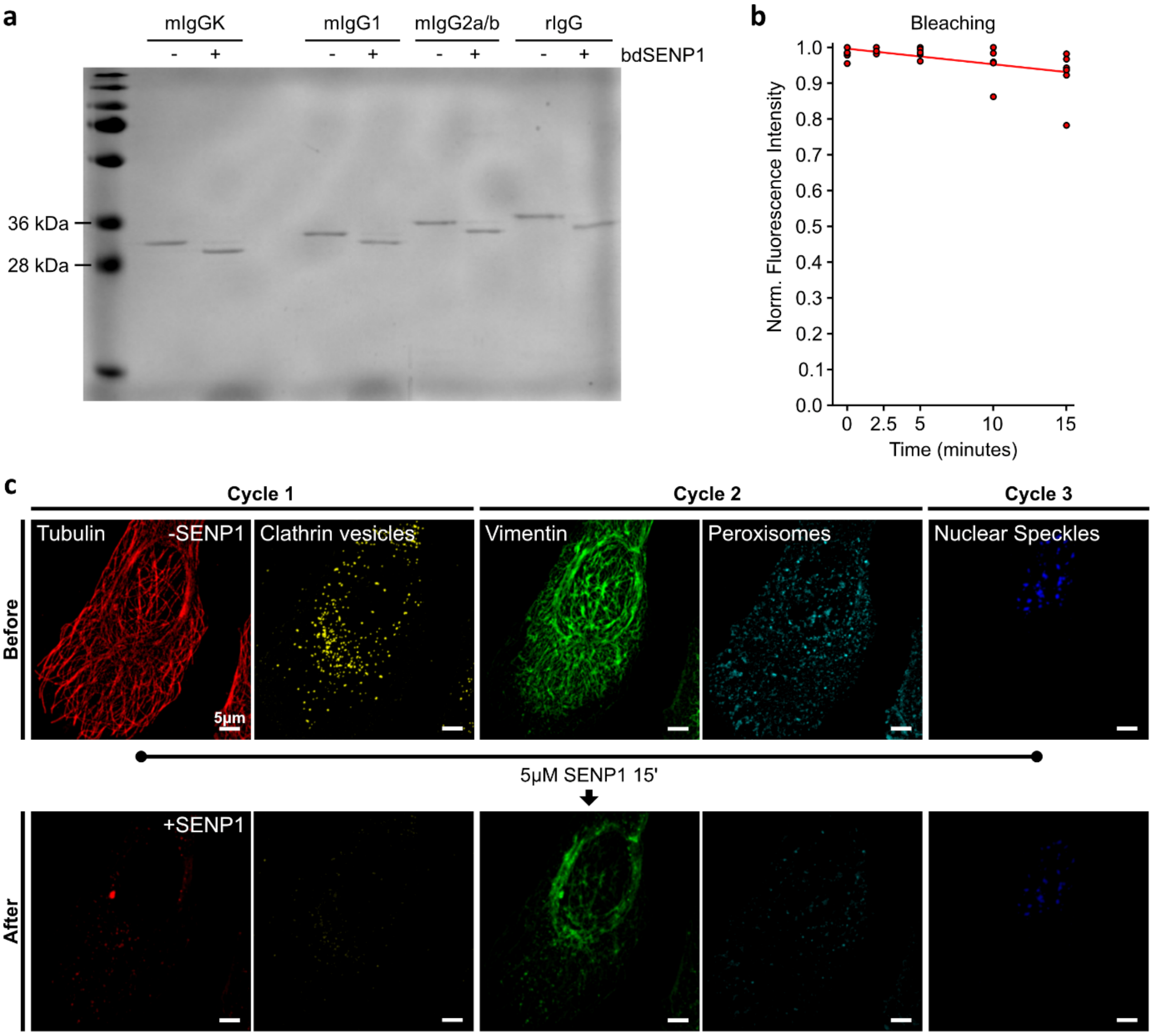


**Supplementary Figure 6. EnzyPlex. a.** Coomassie staining of SDS-PAGE of Enzy-2.Nb anti-Kappa light chain (mIgGK), anti-mouse IgG1 (mIgG1), anti-mouse IgG2a/b (mIgG2a/b) and anti-rabbit IgG (rIgG) incubated with (+) or without (-) 1µM of bdSENP1 for 10 minutes at RT. A molecular weight drop of ~2 KDa can be observed after the Enzy-2.Nbs are exposed to bdSENP1. **b,** Bleaching assessment on six independent samples staining with cleavable 2.Nbs conjugated to Atto643. **c,** Full field of view images from 6 targets during EnzyPlex (Fig. 2) and the corresponding images after cleaving using 5mM bdSENP1 for 15 minutes. Tubulin, clathrin, vimentin, peroxisomes, and nuclear speckles were imaged before and after SENP1 treatment using the same confocal settings. Images are displayed with equal gray levels for direct comparison.

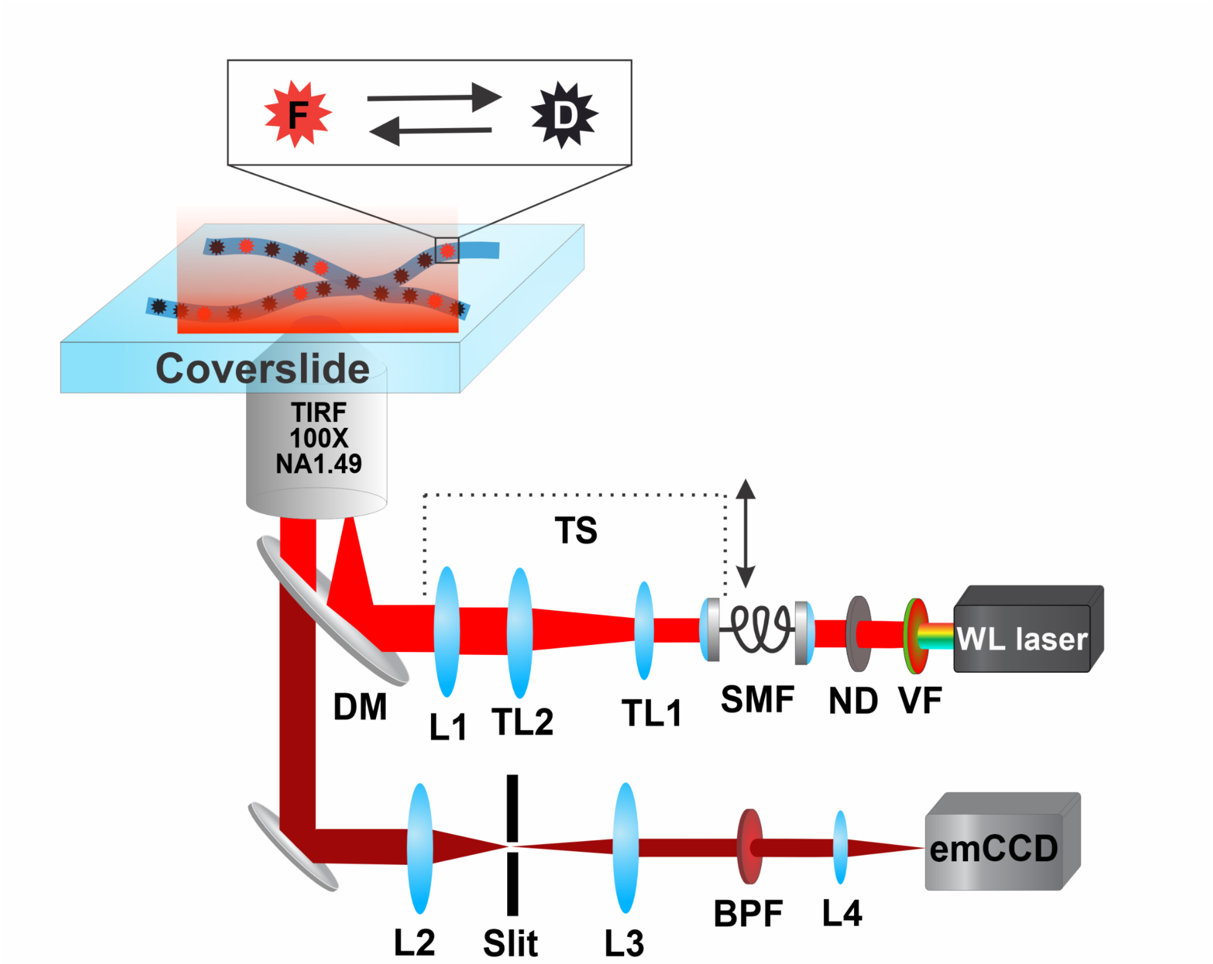


**Supplementary Figure 7: Wide-field SMLM optical setup.** Schematic representation of the custom-built wide-field optical setup for dSTORM imaging. A pulsed super-continuum white light laser (WL Laser) (SuperK Fianium, NKT Photonics) was used for excitation. Variable filter (VF) (SuperK Varia, NKT Photonics) was connected to the laser output and enabled flexibility in the selection of output light wavelength. A neutral density filter (NE10A-A Thorlabs) in tandem with the variable neutral density filter (ND) (NDC-50C-4-A, Thorlabs) were used to adjust the laser excitation power. The laser beam was coupled into a single-mode optical fiber (SMF) (P1-460B-FC-2, Thorlabs). After exiting the optical fiber, the collimated laser beam was expanded by a factor of 3.6X using telescope lenses (TL1 and TL2). The laser beam was focused onto the back focal plane of a TIRF objective (UAPON 100X oil, 1.49NA, Olympus) using an achromatic lens (L1) (AC508-180-AB, Thorlabs). Mechanical shifting of the beam with respect to the optical axis was done through a translation stage (TS) (LNR25/M, Thorlabs) for switching between EPI, HILO, and TIR illumination schemes. The spectral separation of fluorescence light from the excitation pathway was achieved using a multi-band dichroic mirror (DM) (Di03 R405/488/532/635, Semrock), directing the light towards the tube lens (L2) (AC254-200-A-ML, Thorlabs). The field of view was physically limited in the emission path by an adjustable slit aperture (SP60, OWIS) positioned in the image plane. Lenses L3 (AC254-100-A, Thorlabs) and L4 (AC508-150-A-ML, Thorlabs) re-imaged the emitted fluorescence light from the slit onto an emCCD camera (iXon Ultra 897, Andor). A band-pass filter (BP) (BrightLine HC 692/40) was used to block the scattered excitation light further. The total magnification of the optical system on the emCCD camera was 166.6X, resulting in an effective pixel size in the sample space of 103.5nm.

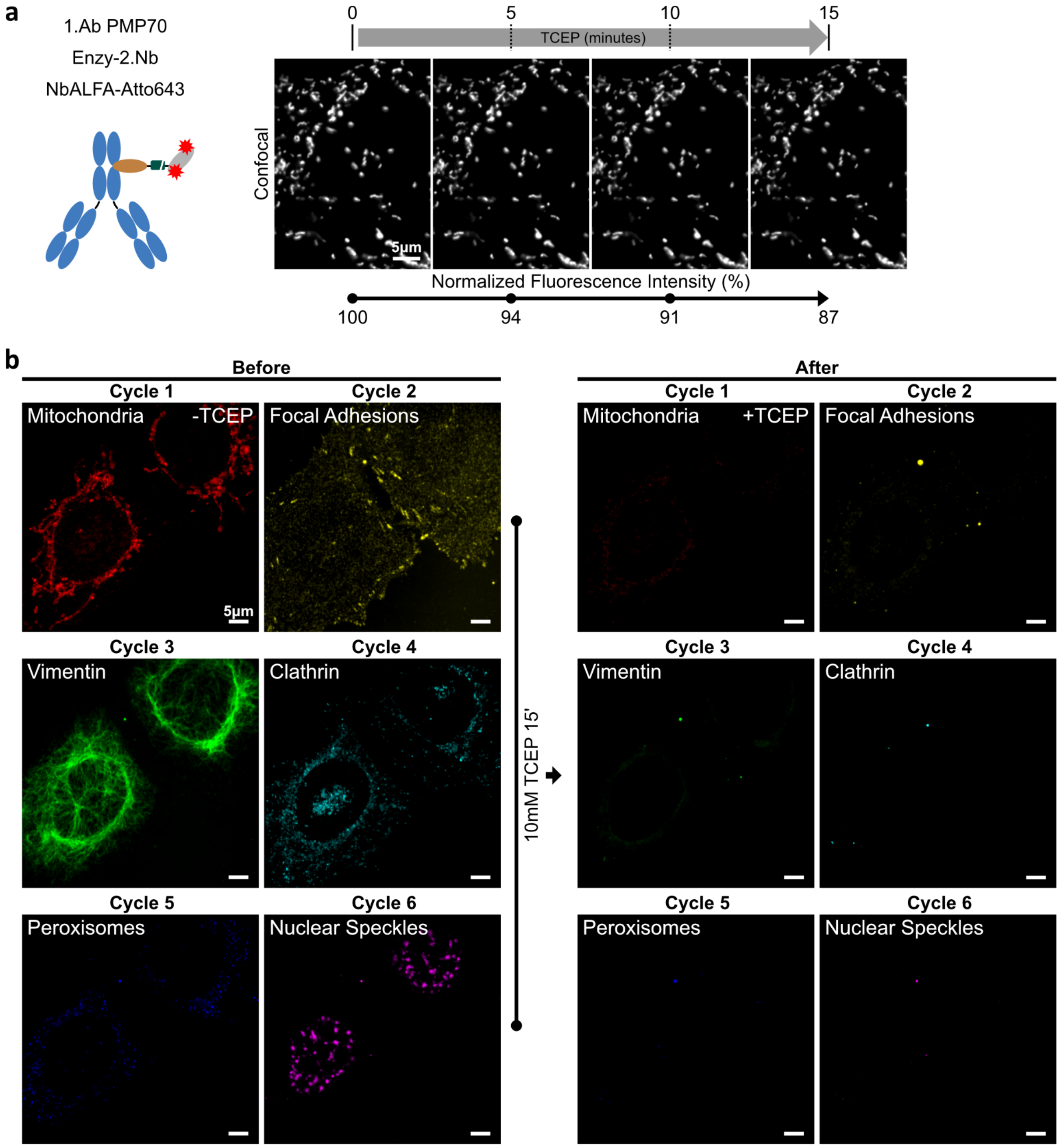


**Supplementary Figure 8. ChemiPlex cleaving efficiency.** **a**, Confocal images of COS-7 cells stained against peroxisomes (1.Ab PMP70) using one-step IF performed with Enzy-2.Nbs. The fluorescence intensity was recorded before treatment with 10mM TCEP-Buffer and at 5, 10, and 15 minutes after treatment. **b,** Full field of view of confocal images in Fig. 3. and the corresponding images after cleaving using 10mM TCEP. Images before and after TCEP treatment were acquired using the same Confocal settings and displayed with equal gray levels for direct comparison.

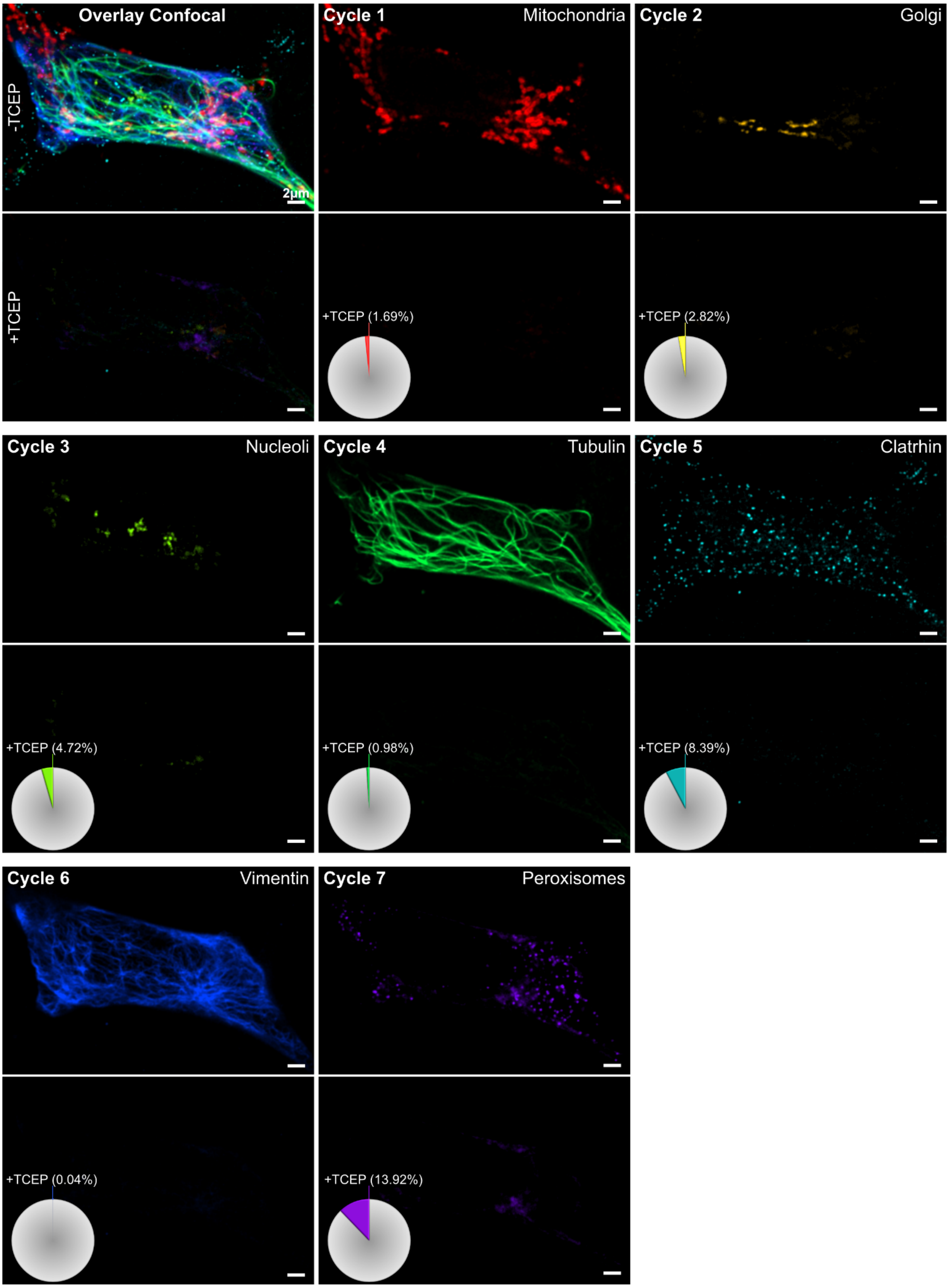


**Supplementary Figure 9. Cleaving efficiency during 8-ChemiPlex STED.** The top-left image displays the overlay of 7 targets in confocal mode, below it, the same overlay of confocal images after the treatment with 10mM TCEP (+TCEP). The same pattern is followed for each of the 7 individual targets (Mitochondria, Golgi, Nucleoli, Tubulin, Clathrin, Vimentin, and Peroxisomes). Images were acquired before and after treatment using the same microscopy settings and are displayed with equally scaled gray values for direct comparison. The pie chart indicates the percentage of the remaining signal in relation to the signal before TCEP cleavage.

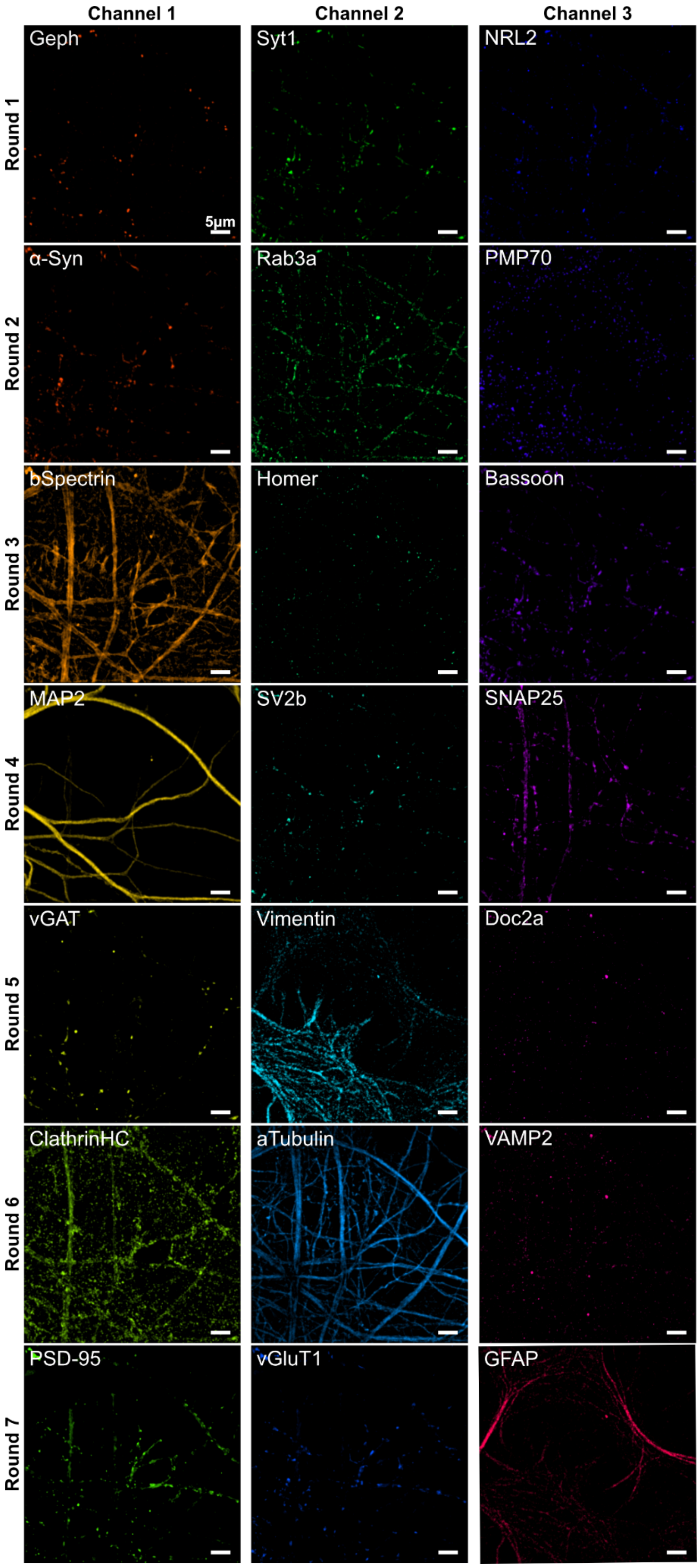


**Supplementary Figure 10. Full field of view of the 21 targets.** Complete volumetric max projections of the 21 confocal images displayed in Fig. 5.

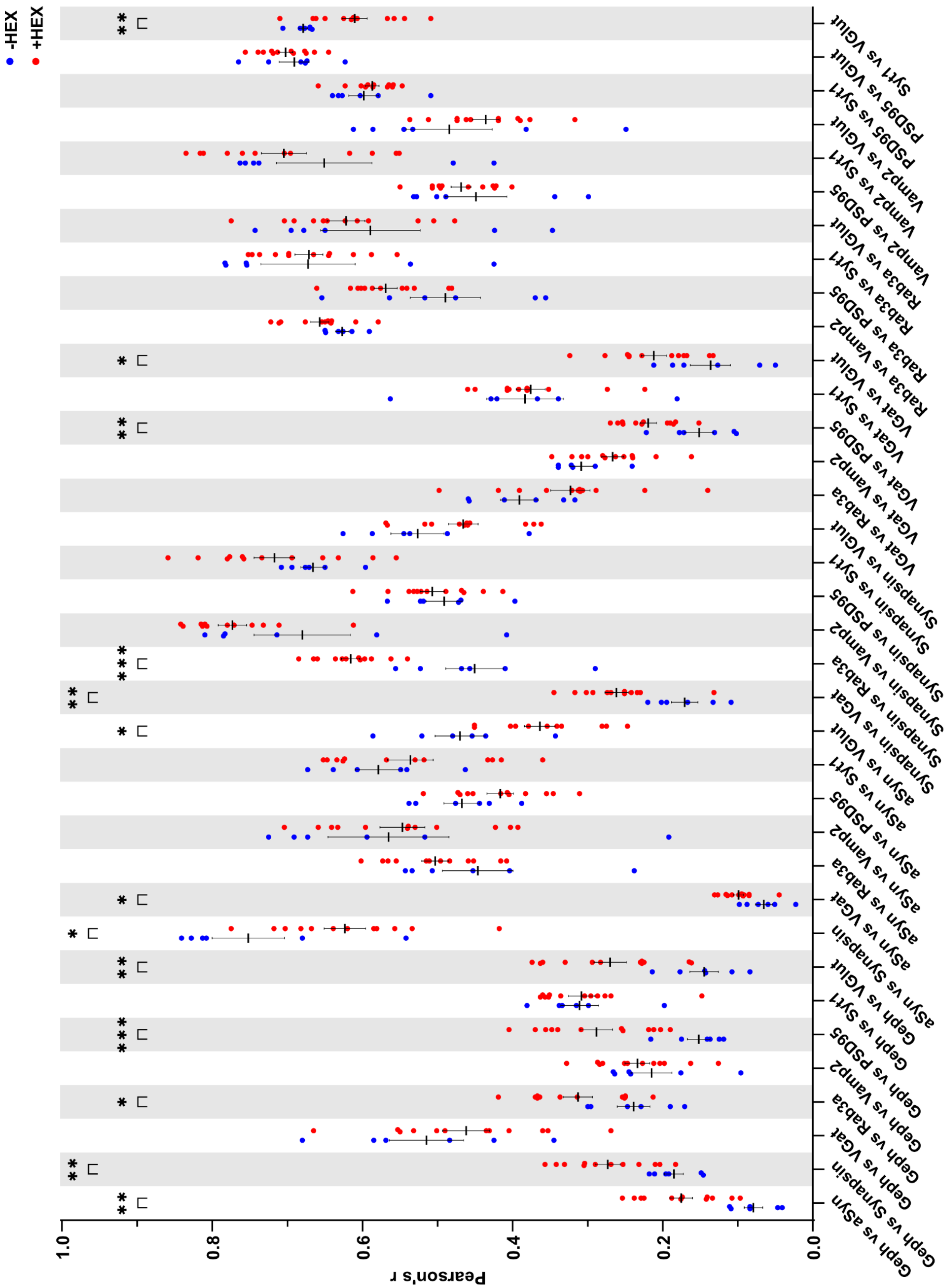

**Supplementary Figure 11. Pearson's correlation between nine synaptic targets with and without 1.6-Hexanodiol.** All the combinations from the 9 targets were compared in correlation before (blue) and after (red) 1.6-Hexanediol treatment. For non-treated samples (-HEX), bars represent the mean value from 6 independent measurements (n = 6). For treated samples (+HEX), bars represent the mean value from 12 independent measurements (n = 12). Error bars show the standard error of the mean (SEM) after unpaired non-parametric Mann-Whitney tests. ∗p<0.05, ∗∗p<0.03, ∗∗∗p<0.002.

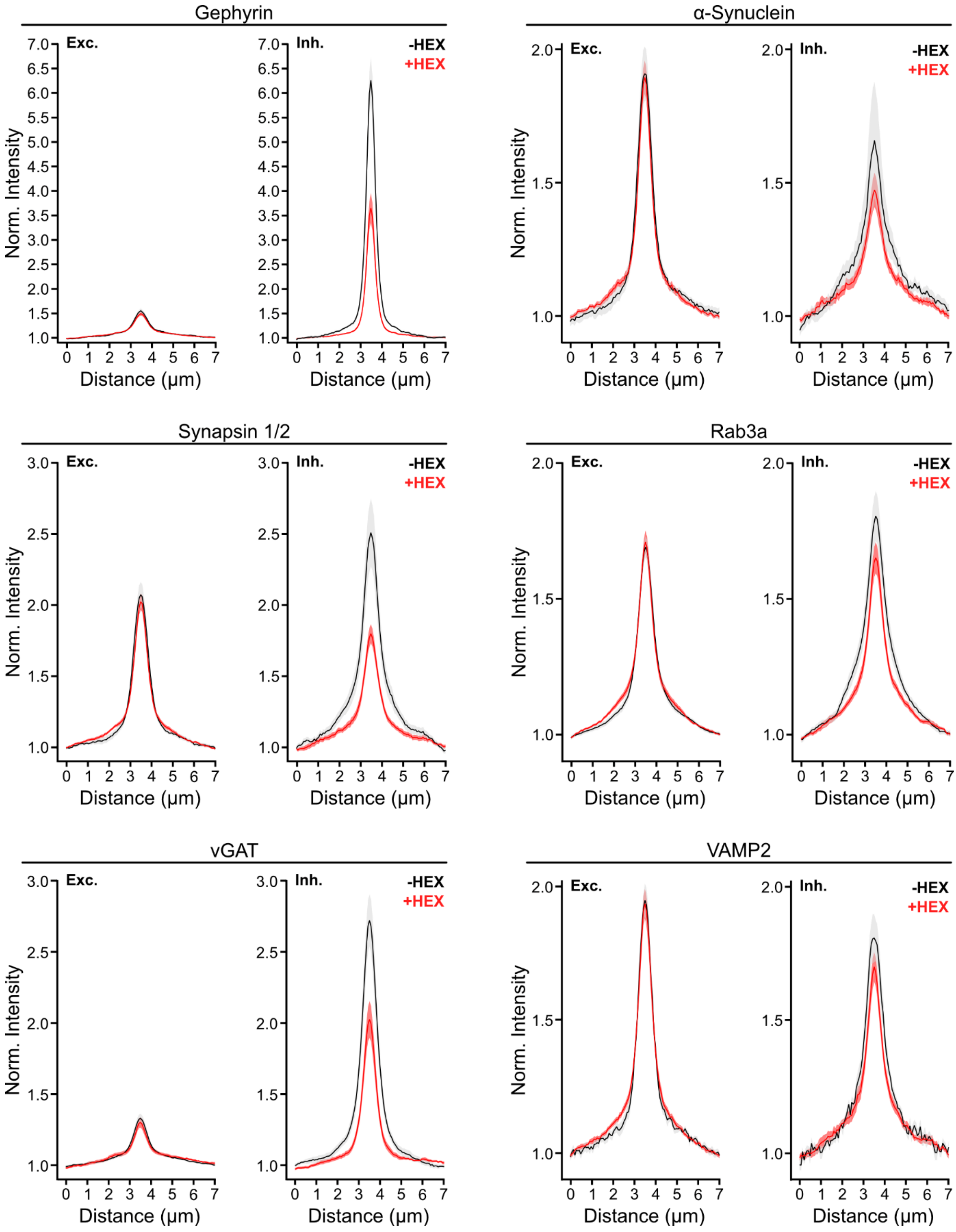


**Supplementary Figure 12. Intensity variation of proteins in excitatory and inhibitory neuronal compartments.** The fluorescence intensity variation of gephyrin, α-Synuclein, Synapsin 1/2, Rab3a, vGAT, and VAMP2 before (-HEX, black) and after (+HEX, red) treatment with 1,6-Hexanediol, measured along excitatory (Exc. PSD-95 positive) and inhibitory (Inh. gephyrin positive) regions of interest. The curves indicate mean ± SEM from n = 10 (-HEX), including 1573 gephyrin-positive and 5663 PSD95-positive synapses; and n = 12 for +HEX treatment, including 3148 gephyrin-positive and 7225 PSD95-positive synapses.

**Supplementary Table 1:** Solvent signals of Chloroform-d1, dimethyl sulfoxide-d6, and methanol-d4.

| **Solvent** | **1H-NMR** | **13C-NMR** |
|---|---|---|
| Chloroform-d3 (CDCl3) | 7.26 ppm | 77.2 ppm |
| Dimethyl sulfoxide-d6 (DMSO-) | 2.50 ppm | 39.5 ppm |
| Methanol-d4 (CD3OD) | 3.31 ppm | 49.0 ppm |

**Supplementary Table 2**: Buffers for NanoPlex

| **Buffer** | **Composition** | **Application** |
|---|---|---|
| **Permeabilization** | 3% BSA<br>0.25% Triton-X100<br>1x PBS | Immunostaining |
| **Washing-Buffer** | 0.1 M Glycine<br>1x PBS<br>pH: 7.5 | Chemi-Plex<br>Enzy-Plex<br>Opto-Plex |
| **Imaging-Buffer** | 1mM PCA<br>1.25 mM Trolox<br>1x PCD (100x PCD: 0.7 mg/ml)<br>1x PBS<br>pH: 7 | Chemi-Plex<br>Enzy-Plex (w/o Trolox) |
| **TCEP-Buffer** | 10 mM TCEP<br>20 mM $K_2CO_3$<br>pH: 6.8 | Chemi-Plex |
| **SENP1-Buffer** | 5 uM SENP1<br>0.1 M Glycine<br>pH: 7 | Enzy-Plex |
| **Thiol-Quenching** | 20 mM NEM<br>0.1 M Glycine<br>0.2 M NaCl<br>1x PBS<br>pH: 7 | Chemi-Plex<br>Enzy-Plex |

**Supplementary Table 3**: Composition of preformed complexes used for Fig. 1f, 1g and the 6-targets OptoPlex immunostainings in U2OS-Nup96-GFP cells on Fig. 1h.

| **Cycle #** | **1.Abs** | **[nM]** | **Opto-2.Nbs** | **[nM]** | **FluoTag®-X2** | **[nM]** |
|---|---|---|---|---|---|---|
| **1** | Alpha-Tubulin | 10 | anti-mouse IgG1 | 30 | anti-ALFA Atto643 | 40 |
| **1** | Clathrin HC | 10 | anti-rabbit IgG | 30 | anti-ALFA AZDye568 | 40 |
| **2** | Vimentin (V9) | 10 | anti-mouse IgG1 | 30 | anti-ALFA Atto643 | 40 |
| **2** | PMP70 | 10 | anti-rabbit IgG | 30 | anti-ALFA AZDye568 | 40 |
| **3** | SON | 15 | anti-rabbit IgG | 45 | anti-ALFA AZDye568 | 60 |
| **3** | - | - | - | - | anti-GFP Atto643 | 20 |

**Supplementary Table 4**: Composition of preformed complexes used for 6-targets EnzyPlex immunostainings in U2OS-Nup96-GFP cells (from Fig. 2).

| Cycle # | 1.Abs | [nM] | Enzy-2.Nbs | [nM] | FluoTag®-X2 | [nM] |
|---|---|---|---|---|---|---|
| 1 | Alpha-Tubulin | 10 | anti-mouse IgG1 | 30 | anti-ALFA Atto643 | 40 |
| 1 | PMP70 | 10 | anti-rabbit IgG | 30 | anti-ALFA AZDye568 | 40 |
| 2 | Vimentin (V9) | 10 | anti-mouse IgG1 | 30 | anti-ALFA Atto643 | 40 |
| 2 | Clathrin HC | 10 | anti-rabbit IgG | 30 | anti-ALFA AZDye568 | 40 |
| 3 | SON | 15 | anti-rabbit IgG | 45 | anti-ALFA AZDye568 | 60 |
| 3 | - | - | - | - | anti-GFP Atto643 | 20 |

**Supplementary Table 5.** Composition of preformed complexes used for 5-targets dSTORM EnzyPlex immunostainings in U2OS-Nup96-GFP cells (from Fig 2).

| Cycle # | 1.Abs | [nM] | Enzy-2.Nbs | [nM] | FluoTag®-X2 | [nM] |
|---|---|---|---|---|---|---|
| 1 | Alpha-Tubulin | 10 | anti-mouse IgG1 | 30 | anti-ALFA JF635b | 40 |
| 2 | Clathrin HC | 10 | anti-rabbit IgG | 30 | anti-ALFA JF635b | 40 |
| 3 | Vimentin (V9) | 10 | anti-mouse IgG1 | 30 | anti-ALFA JF635b | 40 |
| 4 | PMP70 | 10 | anti-rabbit IgG | 30 | anti-ALFA JF635b | 40 |
| 5 | - | - | - | - | anti-GFP JF635b | 20 |

**Supplementary Table 6**: Composition of preformed complexes used for 6-ChemiPlex Confocal in U2OS-Nup96-GFP cells (from Fig. 3).

| Cycle # | 1.Abs | [nM] | Chemi-2.Nbs | [nM] | FluoTag®-X2 | [nM] |
|---|---|---|---|---|---|---|
| 1 | Tom20 (F-10) | 15 | anti-mouse IgG2a/b | 45 | anti-ALFA Atto643 | 60 |
| 2 | Paxillin (Y113) | 15 | anti-rabbit IgG | 45 | anti-ALFA Atto643 | 60 |
| 3 | Vimentin (V9) | 10 | anti-mouse IgG1 | 30 | anti-ALFA Atto643 | 40 |
| 4 | PMP70 | 10 | anti-rabbit IgG | 30 | anti-ALFA Atto643 | 40 |
| 5 | Clathrin HC | 10 | anti-rabbit IgG | 30 | anti-ALFA Atto643 | 40 |
| 6 | - | - | - | - | anti-GFP Atto643 | 20 |

**Supplementary Table 7**: Composition of preformed complexes used for 8-targets STED ChemiPlex immunostainings in U2OS-Nup96-GFP cells (from Fig. 4).

| Cycle # | 1.Abs | [nM] | Chemi-2.Nbs | [nM] | FluoTag®-X2 | [nM] |
|---|---|---|---|---|---|---|
| 1 | Tom20 (F-10) | 15 | anti-mouse IgG2a/b | 45 | anti-ALFA Atto643 | 60 |
| 2 | GALNT2 | 15 | anti-rabbit IgG | 45 | anti-ALFA Atto643 | 60 |
| 3 | UBTF | 15 | anti-rabbit IgG | 45 | anti-ALFA Atto643 | 60 |
| 4 | Alpha-Tubulin | 10 | anti-mouse IgG1 | 30 | anti-ALFA Atto643 | 45 |
| 5 | Clathrin HC | 10 | anti-rabbit IgG | 30 | anti-ALFA Atto643 | 45 |
| 6 | Vimentin (V9) | 10 | anti-mouse IgG1 | 30 | anti-ALFA Atto643 | 45 |
| 7 | PMP70 | 10 | anti-rabbit IgG | 30 | anti-ALFA Atto643 | 45 |
| 8 | - | - | - | - | anti-GFP Atto643 | 20 |

**Supplementary Table 8**: Composition of preformed complexes used for 21-targets ChemiPlex immunostainings in primary hippocampal neurons (from Fig. 5).

| Cycle # | 1.Abs | [nM] | Chemi-2.Nbs | [nM] | FluoTag®-X2 | [nM] |
|---|---|---|---|---|---|---|
| **1** | Gephyrin | 15 | anti-mouse IgG1 | 45 | anti-ALFA Atto643 | 60 |
| **1** | Synaptotagmin1 | 15 | anti-mouse IgG2 | 45 | anti-ALFA AZDye568 | 60 |
| **1** | Neuroligin2 | 15 | anti-mouse IgG1 | 45 | anti-ALFA Atto488 | 60 |
| **2** | Alpha-Synuclein | 15 | anti-mouse IgG1 | 45 | anti-ALFA Atto643 | 60 |
| **2** | Rab3a [15] | 15 | anti-mouse IgG1 | 45 | anti-ALFA AZDye568 | 60 |
| **2** | PMP70 | 10 | anti-rabbit IgG | 30 | anti-ALFA Atto488 | 40 |
| **3** | β-Spectrin II | 15 | anti-mouse IgG1 | 45 | anti-ALFA Atto643 | 60 |
| **3** | Homer1 | 15 | anti-rabbit IgG | 45 | anti-ALFA AZDye568 | 60 |
| **3** | Bassoon | 15 | anti-mouse IgG2 | 45 | anti-ALFA Atto488 | 60 |
| **4** | MAP2 | 15 | anti-mouse IgG1 | 45 | anti-ALFA Atto643 | 60 |
| **4** | SV2b | 15 | anti-mouse IgG2 | 45 | anti-ALFA AZDye568 | 60 |
| **4** | SNAP25 | 15 | anti-mouse IgG1 | 45 | anti-ALFA Atto488 | 60 |
| **5** | VGAT | 15 | anti-rabbit IgG | 45 | anti-ALFA Atto643 | 60 |
| **5** | Vimentin (V9) | 10 | anti-mouse IgG1 | 30 | anti-ALFA AZDye568 | 40 |
| **5** | Doc2a/b | 15 | anti-rabbit IgG | 45 | anti-ALFA Atto488 | 60 |
| **6** | Clathrin HC | 10 | anti-rabbit IgG | 30 | anti-ALFA Atto643 | 40 |
| **6** | Alpha-Tubulin | 10 | anti-mouse IgG1 | 30 | anti-ALFA AZDye568 | 40 |
| **6** | Synaptobrevin2 | 15 | anti-mouse IgG1 | 45 | anti-ALFA Atto488 | 60 |
| **7** | - | | - | | anti-PSD95 Atto643 | 20 |
| **7** | - | | - | | anti-VGlut1 AZDye568 | 20 |
| **7** | - | | - | | anti-GFAP Atto488 | 20 |

**Supplementary Table 9**: Composition of preformed complexes used for 9-targets ChemiPlex immunostainings in primary hippocampal neurons. (from Fig. 6)

| Cycle # | 1.Abs | [nM] | Chemi-2.Nbs | [nM] | FluoTag®-X2 | [nM] |
|---|---|---|---|---|---|---|
| **1** | Gephyrin | 15 | anti-mouse IgG1 | 45 | anti-ALFA Atto643 | 60 |
| **1** | Alpha Synuclein | 15 | anti-mouse IgG1 | 45 | anti-ALFA AZDye568 | 60 |
| **1** | Synapsin1/2 | 15 | anti-rabbit IgG | 45 | anti-ALFA Atto488 | 60 |
| **2** | VGAT | 15 | anti-rabbit IgG | 45 | anti-ALFA Atto643 | 60 |
| **2** | Rab3a | 15 | anti-mouse IgG1 | 45 | anti-ALFA AZDye568 | 60 |
| **2** | Synaptobrevin2 | 15 | anti-mouse IgG1 | 45 | anti-ALFA Atto488 | 60 |
| **3** | - | - | - | - | anti-PSD95 Atto643 | 20 |
| **3** | - | - | - | - | anti-Syt1 AZDye568 | 20 |
| **3** | - | - | - | - | anti-VGlut1 Atto488 | 20 |

**Supplementary Table 10**: Primary antibodies (1.Abs) used in this study.

| Target | Species, Isotype | Company | Cat. # | RRID |
|---|---|---|---|---|
| **Alpha Synuclein** | Mouse, IgG1 | Synaptic Systems | 128 211 | AB_2619811 |
| **Alpha-Tubulin** | Mouse, IgG1 | Synaptic Systems | 30 2211 | AB_887859 |
| **Bassoon** | Mouse, IgG2a | Enzo | ADI-VAM-PS003-F | AB_11181058 |
| **Clathrin** | Rabbit, IgG | Abcam | ab21679 | AB_2083165 |
| **Doc2a/b** | Rabbit, IgG | Synaptic Systems | 174 203 | AB_11064600 |
| **GALNT2** | Rabbit, IgG | Sigma | HPA011222 | AB_1849446 |
| **Gephyrin** | Mouse, IgG1 | Synaptic Systems | 147 011 | AB_887717 |
| **Homer1** | Rabbit, IgG | Synaptic Systems | 160 003 | AB_887730 |
| **MAP2** | Mouse, IgG1 | Synaptic Systems | 188 011 | AB_2147096 |
| **Neuroligin2** | Mouse, IgG1 | Synaptic Systems | 129 511 | AB_2619813 |
| **Paxillin [Y113]** | Rabbit, IgG | Abcam | ab32084 | AB_779033 |
| **PMP70** | Rabbit, IgG | Abcam | ab85550 | AB_10672335 |
| **Rab3a** | Mouse, IgG1 | Synaptic Systems | 107 111 | AB_887770 |
| **SNAP25** | Mouse, IgG1 | Synaptic Systems | 111 111 | AB_887792 |
| **SON** | Rabbit, IgG | Sigma | HPA023535 | AB_1857362 |
| **SV2b** | Mouse, IgG2b | Synaptic Systems | 119 111 | AB_11042616 |
| **Synapsin1/2** | Rabbit, IgG | Synaptic Systems | 106 003 | AB_2619773 |
| **Synaptobrevin2** | Mouse, IgG1 | Synaptic Systems | 104 211 | AB_887811 |
| **Synaptotagmin1** | Mouse, IgG2a | Synaptic Systems | 105 011 | AB_887832 |
| **Tom20 (F-10)** | Mouse, IgG2a | Santa Cruz | sc-17764 | AB_628381 |
| **UBTF** | Rabbit, IgG | Sigma | HPA006385 | AB_1080447 |
| **VGAT** | Rabbit, IgG | ThermoFischer | PA5-27569 | AB_2545045 |
| **Vimentin (V9)** | Mouse, IgG1 | Santa Cruz | sc-6260 | AB_628437 |
| **β-Spectrin II** | Mouse, IgG1 | BD Biosciences | 612562 | AB_399853 |

**Supplementary Table 11**: Nanobodies (Nbs) used in this study.

| Target | Conjugation | Company | Cat. # | RRID |
|---|---|---|---|---|
| **FluoTag®-X2 anti-ALFA** | Atto643 | NanoTag | N1502-At643-L | AB_3075983 |
| **FluoTag®-X2 anti-ALFA** | AZDye568 (AF568) | NanoTag | N1502-AF568-L | AB_3075980 |
| **FluoTag®-X2 anti-ALFA** | Atto488 | NanoTag | N1502-At488-L | AB_3075982 |
| **FluoTag®-X2 anti-ALFA** | Janelia Fluor635b (JF635b) | NanoTag | Custom made | NA |
| **FluoTag®-X2 anti-GFAP** | Atto488 | NanoTag | N3802-At488-L | AB_3076117 |
| **FluoTag®-X2 anti-PSD95** | Atto643 | NanoTag | N3702-At643-L | AB_3076106 |
| **FluoTag®-X2 anti-Syt1 (A51)** | AZDye568 (AF568) | NanoTag | N4302-AF568-L | AB_3076136 |
| **FluoTag®-X2 anti-VGlut1** | AZDye568 (AF568) | NanoTag | N1602-AF568-L | AB_3076003 |
| **FluoTag®-X2 anti-VGlut1** | Atto488 | NanoTag | N1602-At488-L | AB_3076005 |
| **FluoTag®-X4 anti-GFP** | Atto643 | NanoTag | N0304-At643-L | AB_3075906 |
| **Multiplexing Blocker Mouse** | w/o | NanoTag | K0102-50 | NA |
| **Multiplexing Blocker Rabbit** | w/o | NanoTag | K0202-50 | NA |
| **sdAb anti-ALFA** | Unconj. C-term. Cys | NanoTag | N1505-250ug | AB_3075986 |
| **sdAb anti-Mouse IgG1** | Unconj. C-term. Cys | NanoTag | N2005-250ug | AB_3076025 |
| **sdAb anti-Mouse IgG2a/b** | Unconj. C-term. Cys | NanoTag | N2705-250ug | AB_3076057 |
| **sdAb anti-Rabbit IgG** | Unconj. C-term. Cys | NanoTag | N2405-250ug | AB_3076046 |

**Detailed Synthetic Procedures and Compound Characterization**

**Standard Operating Procedures (SOPs)**

*SOP1: Automated Solid Phase Peptide Synthesis (SPPS)*

A Liberty Blue peptide synthesizer from CEM (Matthews, North Carolina, USA) was used to synthesize different peptide sequences. The following protected amino acids were dissolved in DMF: Fmoc-Arg(Pbf)-OH, Fmoc-Glu(O$^t$Bu)-OH, Fmoc-Gly-OH, Fmoc-Leu-OH, Fmoc-Pro-OH, Fmoc-Ser(O$^t$Bu)-OH, Fmoc-Thr(O$^t$Bu)-OH. Depending on the synthesis scale, different coupling conditions, different stock solutions for the amino acids, deprotection solution, activator, and activator base were used in compliance with the recommended CEM protocols for the Liberty Blue Synthesizer. For the deprotection of the Fmoc group, 20% piperidine in DMF (*v/v*) was used. The amino acids were activated by DIC as the activator and Oxyma as the activator base. The non-preloaded resin (1.0 eq) was placed in the reaction vessel of the peptide synthesizer and was swollen for 5 min in DMF. First, the *N*-terminal Fmoc protecting group was removed by the addition of piperidine (20 % in DMF, *v/v*) under microwave irradiation (1: 75 °C, 90 W, 15 s; 2: 90 °C, 20 W, 50 s). To achieve a complete cleavage of the Fmoc-group, the deprotection step was repeated twice. After washing the resin with DMF (5 x 4 mL), the amino acid, DIC, and Oxyma were added to the reaction vessel. The coupling reaction was performed under microwave irradiation (1: 75 °C, 170 W, 15 s; 2: 90 °C, 30 W, 110 s). For the coupling of Fmoc-Arg(Pbf)-OH (75 °C, 30 W, 300 s), recommended coupling cycles were used to suppress γ-lactam formation. After the peptide synthesis was completed, the resin was transferred into a BD Syringe with PE-frit and was washed with DMF (6 x 4 mL) and DCM (6 x 4 mL). The resin was then dried under reduced pressure.

**Supplementary Table 12**: Overview of coupling and deprotection conditions on different synthesis scales.

| | 0.01 mmol | 0.05/0.025 mmol |
|---|---|---|
| Deprotection | 2 % Piperazine | 20 % Piperidine |
| Amino acids | 0.08 M | 0.2 M |
| Activator | 0.05 M DIC | 0.25 M DIC |
| Activator Base | 0.1 M Oxyma | 0.5 M Oxyma |
| Deprotection 1.: | 75 °C, 155 W, 15 s | 75 °C, 210 W, 15 s |
| 2.: | 90 °C, 30 W, 165 s | 90 °C, 30 W, 50 s |
| Coupling 1.: | 75 °C, 170 W, 15 s | 75°C, 170 W, 15 s |
| 2.: | 90 °C, 30 W, 465 sec | 90°C, 30 W, 110 s |

*SOP2: Cleavage of Peptide Strands from Resin*

The cleavage of the peptides from the resin was performed in a BD syringe with a PE frit. The resin was shaken at room temperature for 2 h in a TFA/Thioanisole/EDT/anisole solution (90:5:3:2 v/v/v/v). After cleavage, the solution was concentrated under a nitrogen stream and the crude peptide was precipitated with diethyl ether. The precipitate was isolated by centrifugation (9000 rpm, -10 °C, 10 min), washed twice with ether, and dried under vacuum.

*Experimental Procedures*

**5-methoxy-2-nitro-4-(prop-2-yn-1-yloxy) benzaldehyde (2):** To a solution of 4-hydroxy-5-methoxy-2-nitrobenzaldehyde (200 mg, 1.01 mmol, 1.00 eq,) and $K_2CO_3$ (210 mg, 1.52 mmol, 1.50 eq) in DMF (3 mL), propargyl bromide (174 mg, 1.22 mmol, 1.20 Eq) was added and the mixture was stirred for 16 hours at room temperature. The solution was then washed with water (2 x 10 mL) and extracted with EtOAc (3 x 30 mL). The combined organic phases were dried over $MgSO_4$ and the solvent was removed under reduced pressure. The product (155 mg, 0.66 mmol, 65%) was isolated as a light brownish solid. $^1$H-NMR (300 MHz, $CDCl_3$): $\delta$ = 10.47 (s, 1 H), 7.81 (s, 1 H), 7.45 (s, 1 H), 4.94 (d, 2 H), 4.04 (s, 3 H), 2.66 (m, 1 H). $^{13}$C-NMR (75 MHz, $CDCl_3$): $\delta$ = 187.7, 153.7, 149.9, 143.9, 126.4, 110.3, 109.4, 77.8, 76.4, 57.2, 56.8. HR-MS (ESI): m/z calculated for $C_{11}H_8NO_5[M-H]^-$: 234.0408, found: 234.0414.

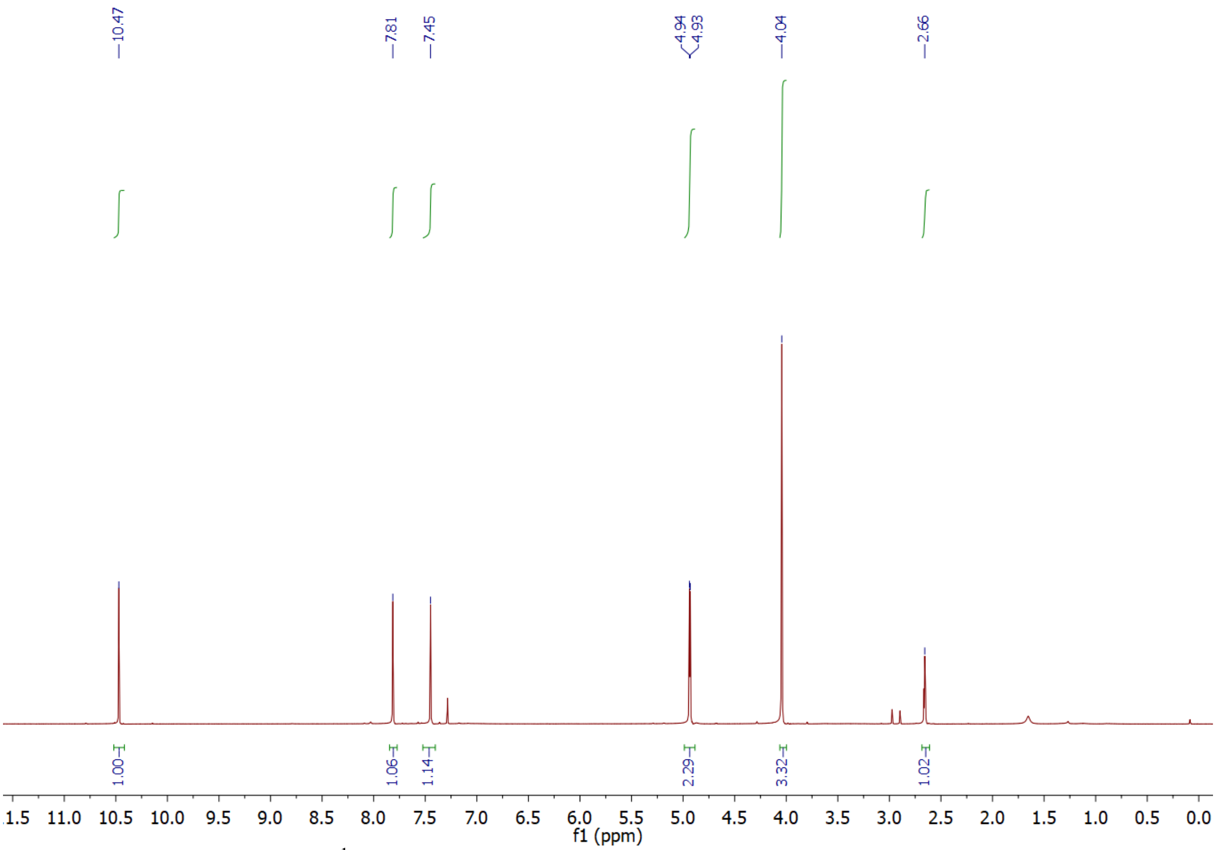


**Supplementary Figure 13:** $^1$H-NMR of compound **2** in $CDCl_3$.

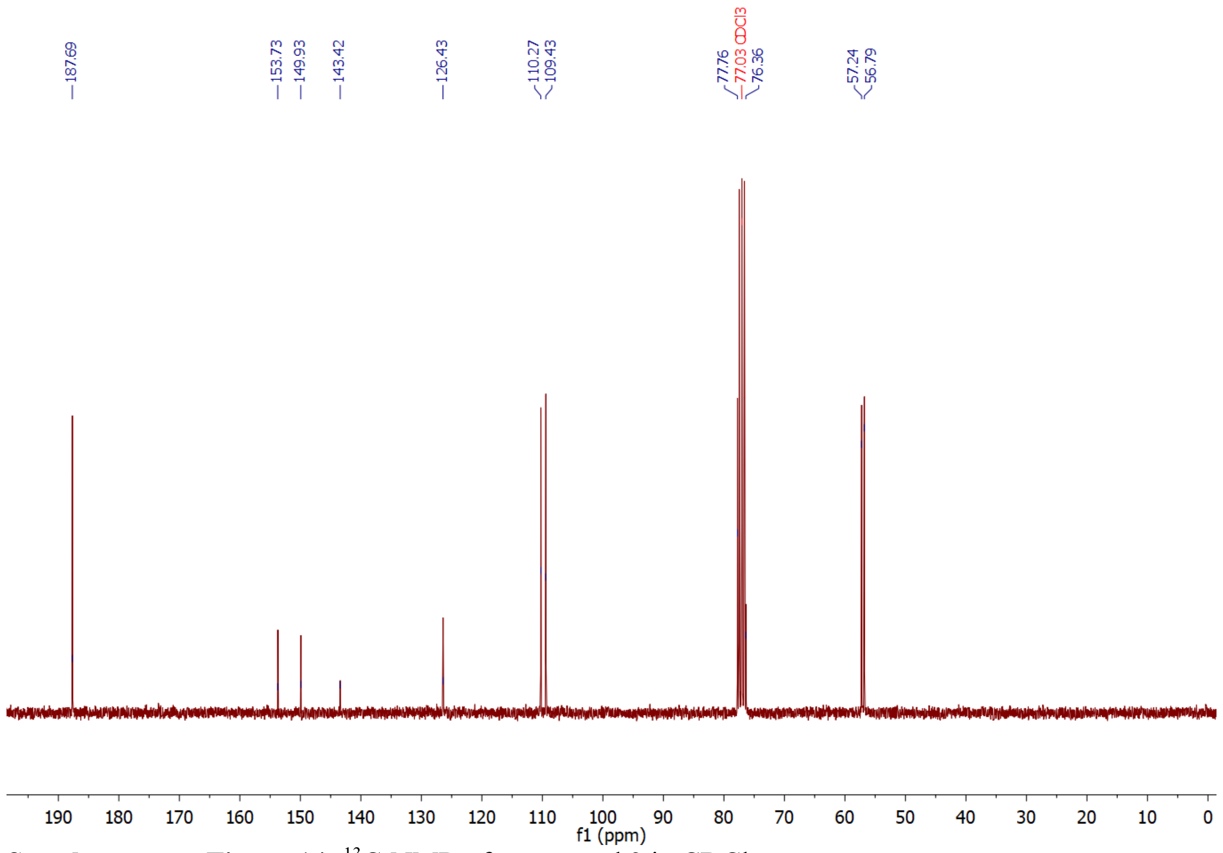


**Supplementary Figure 14:** $^{13}$C-NMR of compound **2** in $CDCl_3$.

**(5-Methoxy-2-nitro-4-(prop-2-yn-1-yloxy)phenyl)methanol (3):** To a solution of 2 (129 mg, 0.55 mmol, 1.00 eq.) in methanol (5 mL) was added $NaBH_4$ (42 mg, 1.10, mmol, 2.00 eq.) and the mixture was stirred for 30 min. The solution was neutralized with 2 M HCl and the solvent was removed under reduced pressure. The residue was dissolved in DCM (1 mL), washed with Water (2 x 10 mL) and then extracted with EtOAc (3 x 20 mL). After the solvent was removed, the product (114 mg, 0.48 mmol, 87%) was isolated as a crystalline solid. $^{1}$H-NMR (300 MHz, $CDCl_3$): $\delta$ = 7.90 (s, 1 H), 7.24 (s, 1 H), 5.00 (s, 1 H), 4.86 (m, 2 H) 4.02 (s, 3 H), 2.66 (s, 1 H), 2.60 (s, 1 H). $^{13}$C-NMR (75 MHz, $CDCl_3$): $\delta$ = 154.5, 145.5, 139.5, 133.4, 111.4, 111.0, 77.2, 77.0, 62.8, 57.1, 56.5. HR-MS (ESI): m/z calculated for $C_{11}H_{12}NO_5[M+H]^+$ : 238.0716, found: 238.0412.

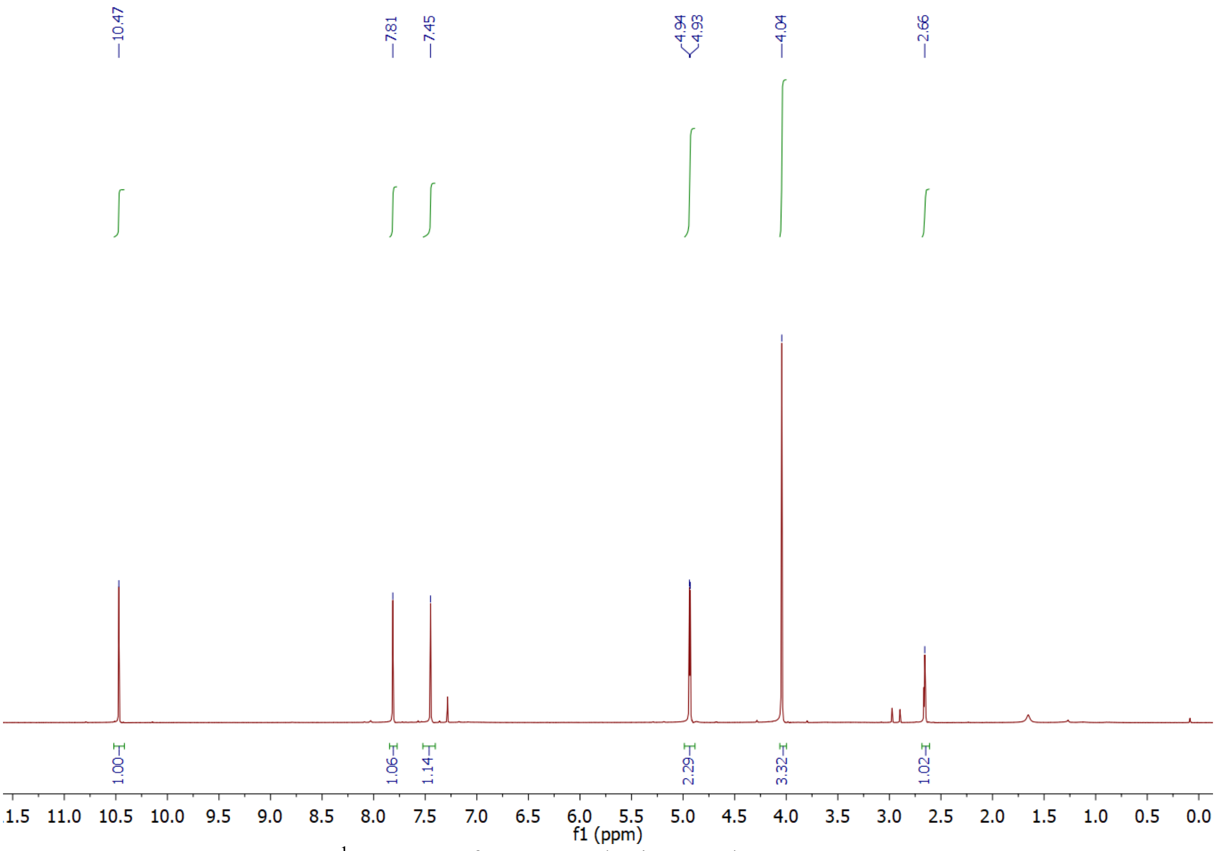


**Supplementary Figure 15:** $^{1}$H-NMR of compound **3** in $CDCl_3$.

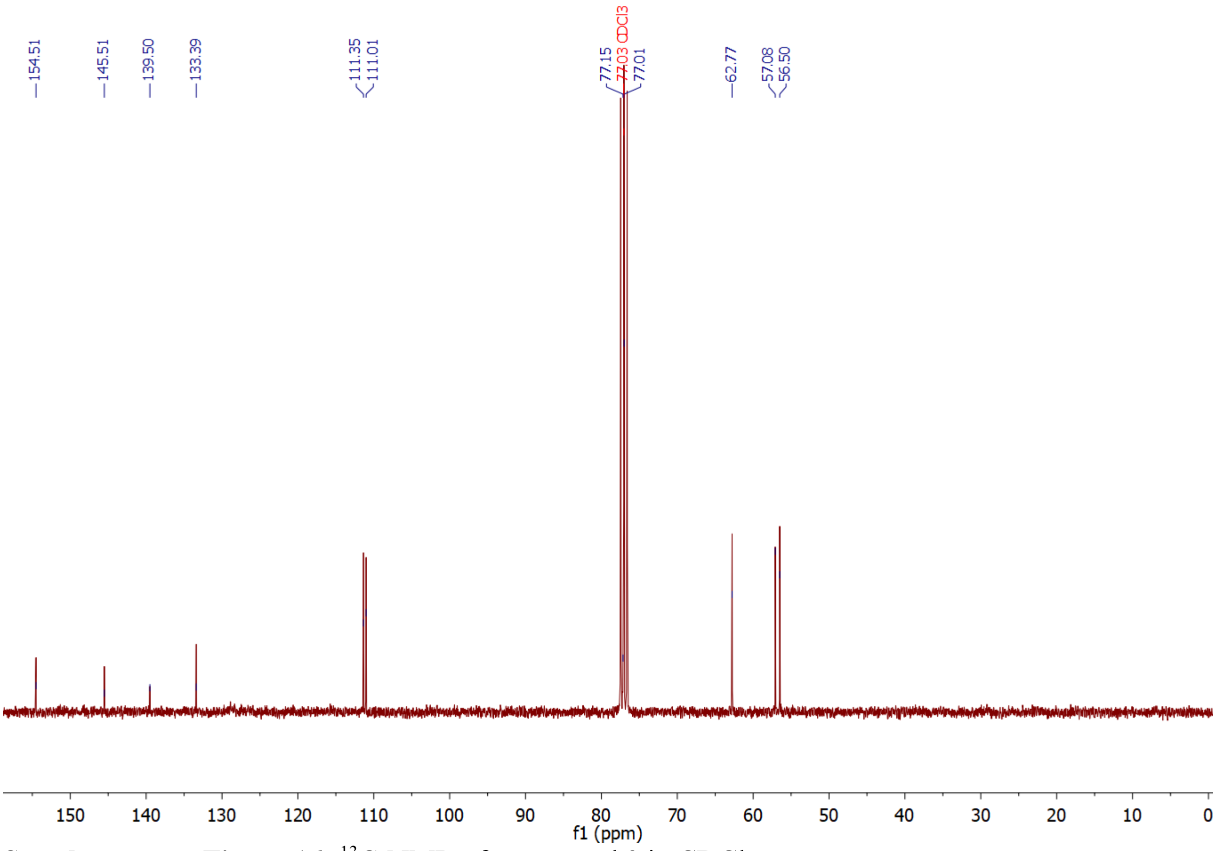


**Supplementary Figure 16:** $^{13}$C-NMR of compound **3** in $CDCl_3$.

**1-(Bromomethyl)-5-methoxy-2-nitro-4-(prop-2-yn-1-yloxy)benzene (4): 3** (732 mg, 3.0 mmol, 1.0 eq.) was dissolved in DCM (25 mL) and chilled to 0°C. Afterwards, $PBr_3$ (2.85 g, 1 mL, 10 mmol, 3.3 eq.) was added slowly and the mixture was stirred for 12 h, while reaching room temperature. The reaction mixture was washed with water (50 mL) and the aqueous phase was extracted with EtOAc (3 x 30 mL). The organic phase was dried over $MgSO_4$ and the solvent was removed *in vacuo*. The final product (863 mg, 2.88 mmol, 95%) was yielded as yellowish solid. $^{1}$H-NMR (300 MHz, $CDCl_3$): $\delta$ = 7.84 (s, 1H), 6.98 (s, 1H), 4.87 (s, 2H), 4.84 (d, $J$ = 2.4 Hz, 2H), 3.99 (s, 3H), 2.59 (t, $J$ = 2.4 Hz, 1H ). $^{13}$C-NMR (75 MHz, $CDCl_3$): $\delta$ =153.9, 146.6, 140.2, 128.6, 114.2, 111.3, 77.4, 77.0, 57.2, 56.7, 30.0. HR-MS (ESI): m/z berechnet für $C_{11}H_{10}NO_4Br$ $[M+Na]^+$: 321.9685, found: 321.9687. IR (ATR) [$cm^{-1}$]: 3281, 2360, 2339, 1521, 1509, 1325, 1270, 1228, 1062, 876. Mp.: 124–127 °C.

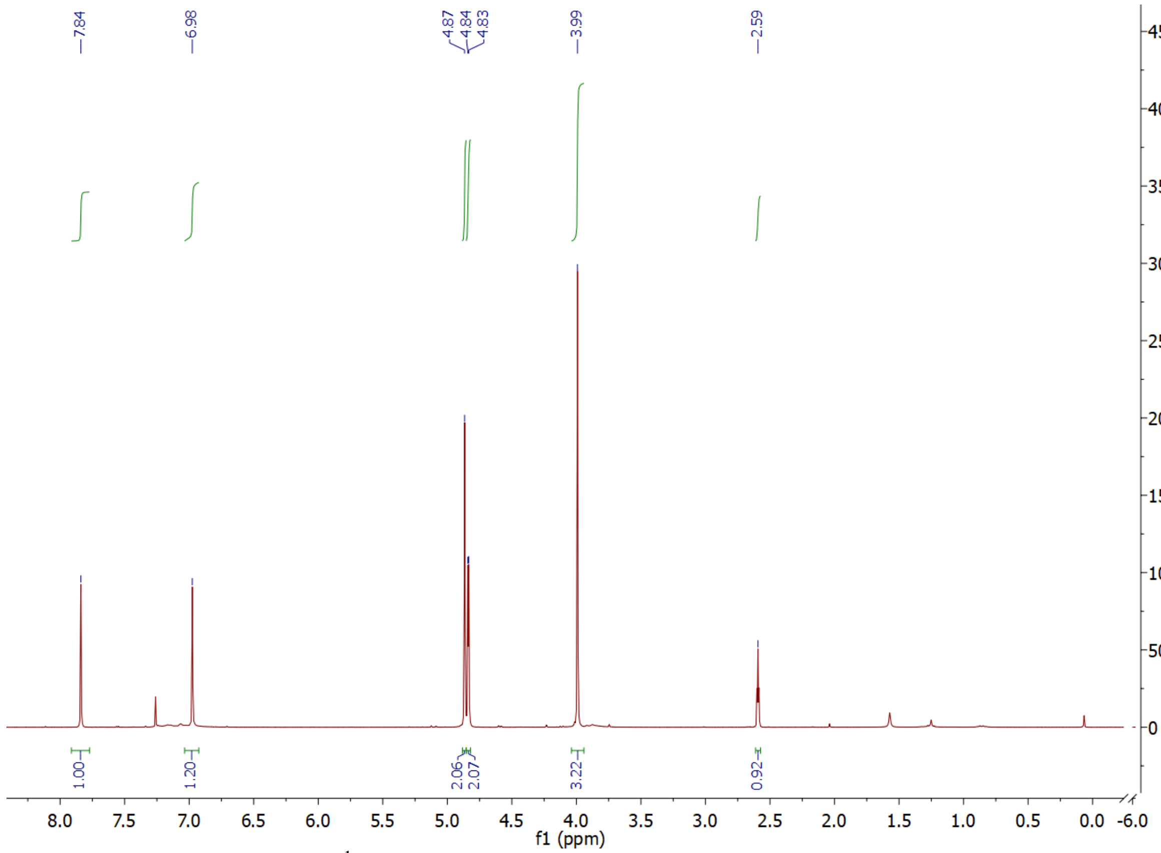


**Supplementary Figure 17:** $^{1}$H-NMR of compound **4** in $CDCl_3$.

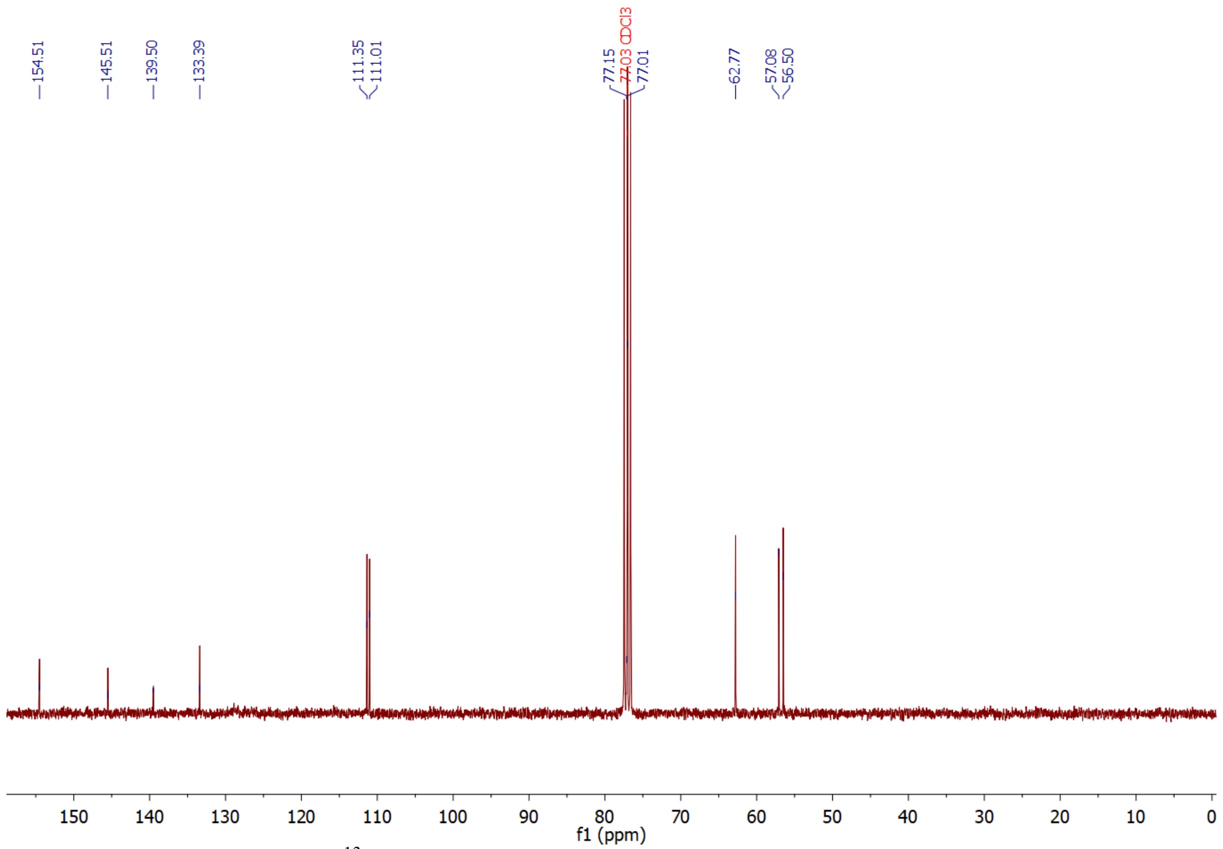


**Supplementary Figure 18:** $^{13}$C-NMR of compound **4** in $CDCl_3$.

**(4-((5-methoxy-2-nitro-4-(prop-2-yn-1-yloxy)benzyl)oxy)phenyl)methanol (6):** Potassium carbonate (1.51 g, 10.9 mmol, 2.0 Eq.), 4-hydroxybenzyl alcohol (690 mg, 5.56 mmol, 1.02 Eq.) and 4 (1.63 g, 5.46 mmol, 1.0 Eq.) were added to acetone (55 mL) and refluxed for 3 h. The solvent was afterward removed, water (50 mL) was added and the mixture was extracted with EtOAc (3 x 50 mL). The organic phase was dried, and the solvent was removed *in vacuo* to yield the crude product, which was purified by column chromatography (20% EtOAc/ Pentane) on silica to yield the final compound (1.96 g, 5.42 mmol, 99%) as yellowish crystals. $^{1}$H-NMR (300 MHz, $CDCl_3$): $\delta$ = 7.94 (s, 1H), 7.37 (s, 1H), 7.32 (d, $J$ = 8.5 Hz, 1H), 6.99 (d, $J$ = 8.5 Hz, 1H), 5.50 (s, 2H), 4.84 (d, $J$ = 2.4 Hz, 2H), 4.64 (s, 2H), 3.95 (s, 3H), 2.59 (t, $J$ = 2.4 Hz, 1H). $^{13}$C-NMR (101 MHz, $CDCl_3$): δ 157.7, 154.6, 145.4, 138.8, 134.1, 130.5, 128.8, 115.1, 110.8, 109.9, 77.3, 77.2, 67.2, 64.9, 57.1, 56.5. HR-MS (ESI): m/z calculated for $C_{18}H_{17}NO_6$ $[M+Na]^+$: 366.0948, found: 366.0951. IR (ATR) [$cm^{-1}$]: 2360, 2339, 1512, 1332, 1273, 1240, 1210, 1179, 1062, 992, 980. Mp.: 149–151 °C.

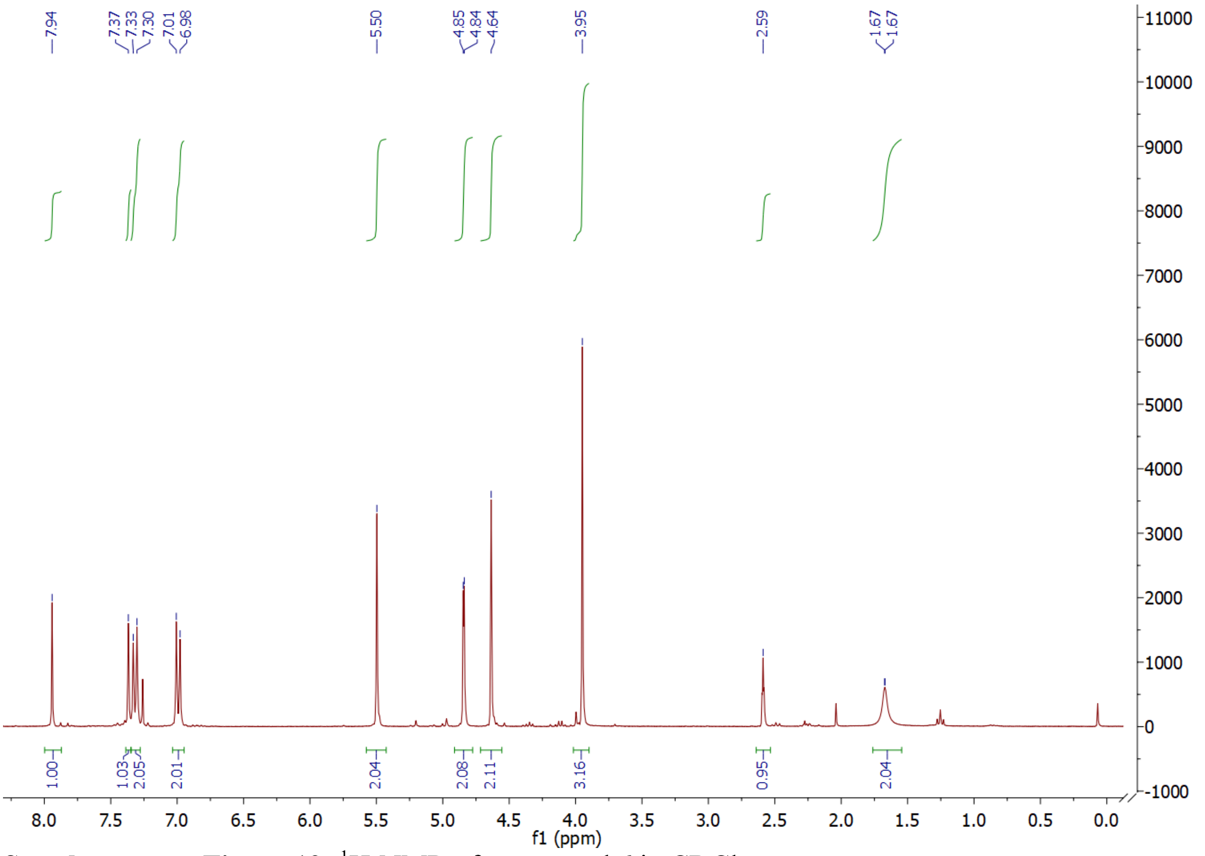


**Supplementary Figure 19:** $^{1}$H-NMR of compound **6** in $CDCl_3$.

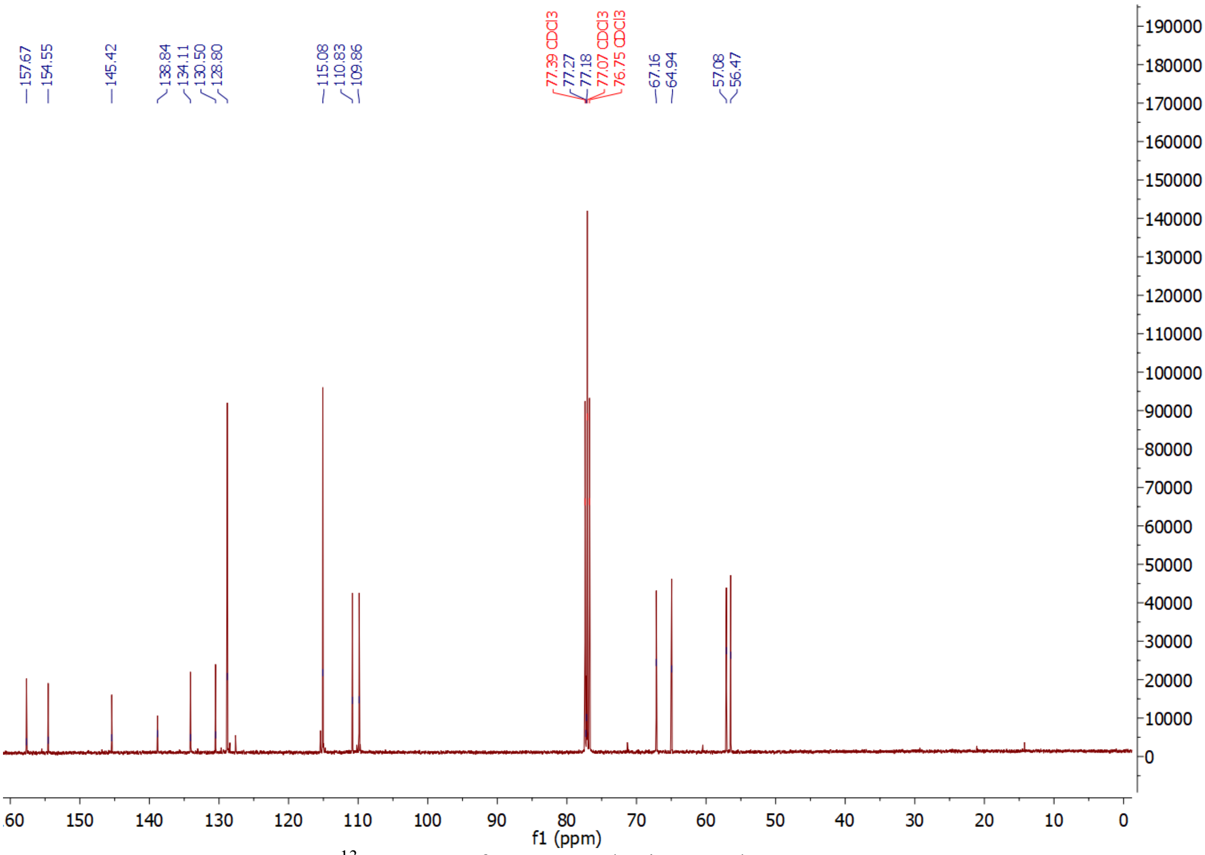


**Supplementary Figure 20:** $^{13}$C-NMR of compound **6** in $CDCl_3$.

**1-((4-(Bromomethyl)phenoxy)methyl)-5-methoxy-2-nitro-4-(prop-2-yn-1-yloxy)benzene (7):** 6 (1.50 g, 4.4 mmol, 1.0 Eq.) was dissolved in DCM (80 mL) and the reaction mixture was chilled with an ice bath. Afterwards, $PBr_3$ (1.4 mL, 15.1 mmol, 3.4 Eq.) was added slowly and the mixture was allowed to slowly reach room temperature and was stirred at this temperature for 2d. Water (50 mL) was added, the organic phase was separated and the aqueous phase was extracted with EtOAc (3 x 50mL). The combined organic phases were dried with magnesium sulfate and the solvent was removed in vacuo to yield the crude product, which was purified by column chromatography (20% EtOAc/Pentane) on silica to yield the final compound (1.16 g, 2.86 mmol, 65 %) as yellowish solid. $^1$H-NMR (300 MHz, $CDCl_3$): $\delta$ =7.94 (s, 1H), 7.36 (s, 1H), 7.32 (d, $J$ = 8.6 Hz, 2H), 6.99 (d, $J$ = 8.7 Hz, 2H), 5.50 (s, 2H), 4.84 (d, $J$ = 2.4 Hz, 2H), 4.63 (s, 2H), 3.95 (s, 3 H), 2.59 (t, $J$ = 2.4 Hz, 1H). $^{13}$C-NMR (75 MHz, $CDCl_3$): $\delta$ = 158.2, 154.7, 145.6, 139.0, 131.1, 130.8, 130.3, 115.4, 111.0, 110.0, 77.3, 67.3, 60.5, 57.2, 56.6, 33.7, 21.2, 14.3. HR-MS (ESI): m/z calculated for $C_{18}H_{16}BrNO_6$ $[M+Na]^+$: 430.0084, found: 430.0073. IR (ATR) [$cm^{-1}$]:3300, 1604, 1586, 1513, 1322, 1271, 1253, 1216, 1069, 1033. Mp.: 134–136 °C.

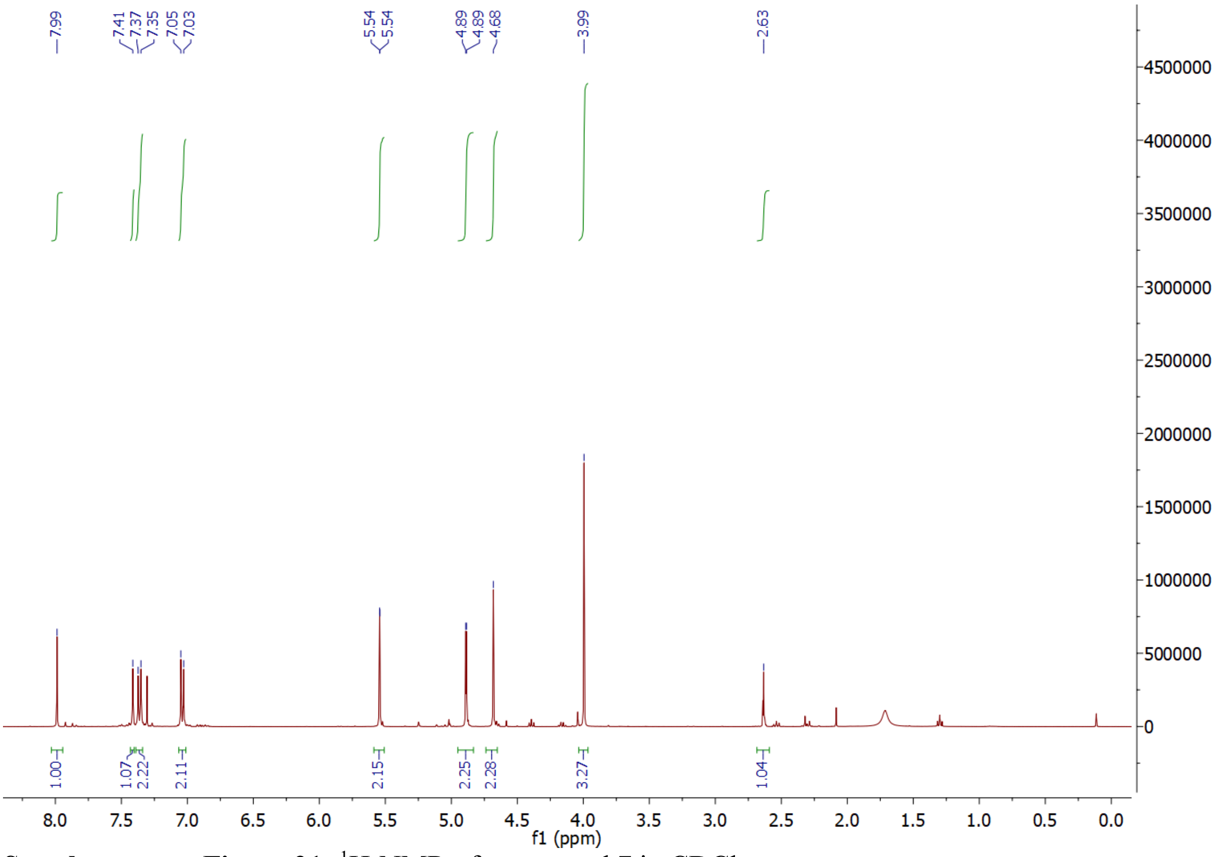


**Supplementary Figure 21:** $^1$H-NMR of compound **7** in $CDCl_3$.

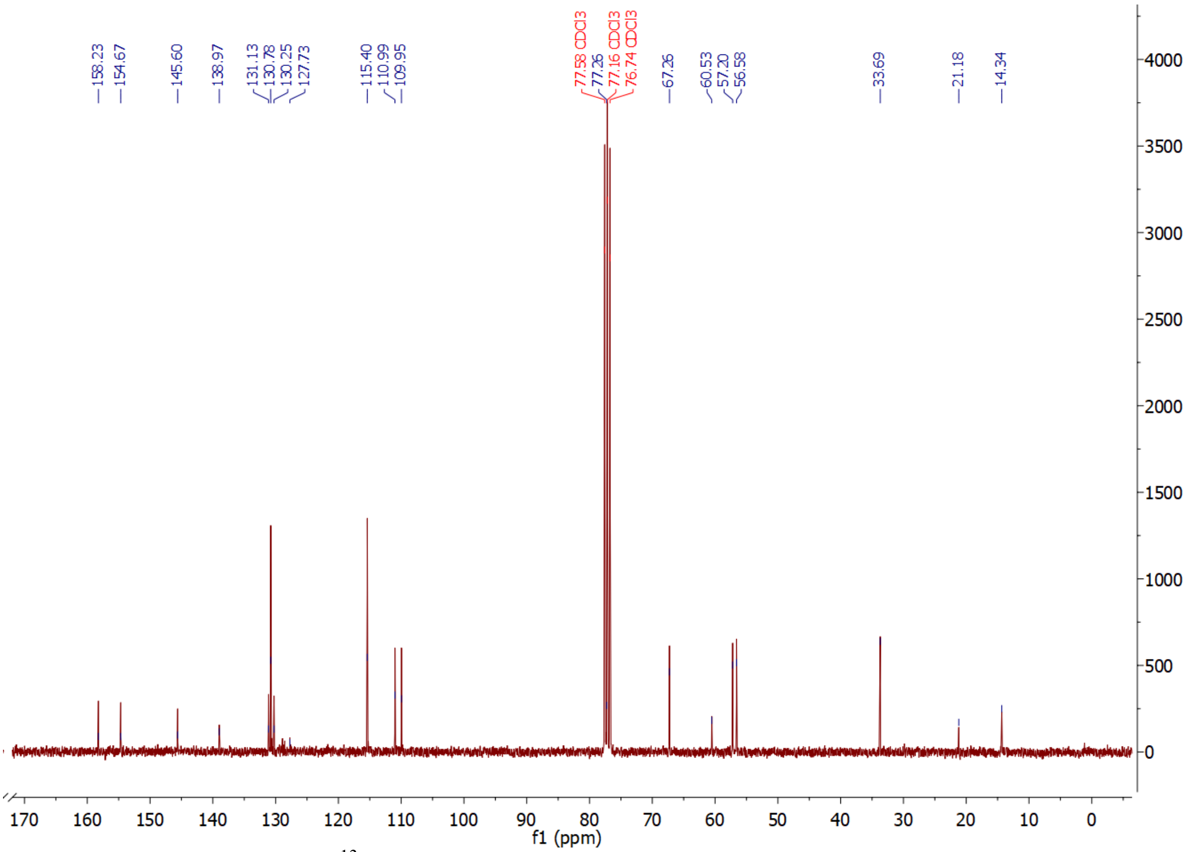


**Supplementary Figure 22:** $^{13}$C-NMR of compound **7** in $CDCl_3$.

**3a,4,7,7a-Tetrahydro-1H-4,7-epoxyisoindole-1,3(2H)-dione (8):** Under inert conditions, maleimide (5.00 g, 51.5 mmol, 1.0 Eq.) was dissolved in dioxane (76 mL), furan (12 mL, 0.154 mol, 3.0 Eq.) was added and the reaction mixture was stirred at 90°C for 12 h. The reaction mixture was poured into water (250 mL) and the aqueous phase was extracted with EtOAc (3 x 200 mL). The organic phase was washed with brine, dried over $MgSO_4$ and the solvent was removed in vacuo the yield the final compound (7.89 g, 47.8 mmol, 92%) as light brownish solid. $^1$H-NMR (300 MHz, DMSO): $\delta$ = 11.15 (s, 1H), 6.53 (s, 2H), 5.12 (s, 2H), 2.85 (s, 2H). $^{13}$C-NMR (75 MHz, DMSO): $\delta$ = 178.3, 137.0, 80.8, 48.9. HR-MS (ESI): m/z calculated for $C_8H_7NO_3$ $[M-H]^-$: 163.0353, found: 163.0353. IR (ATR) [$cm^{-1}$]: 3164, 3067, 2360, 1769, 1698, 1350, 1283, 1203, 1188, 1139, 1087, 1023, 839, 732. Mp.: 153–155 °C (decomposition).

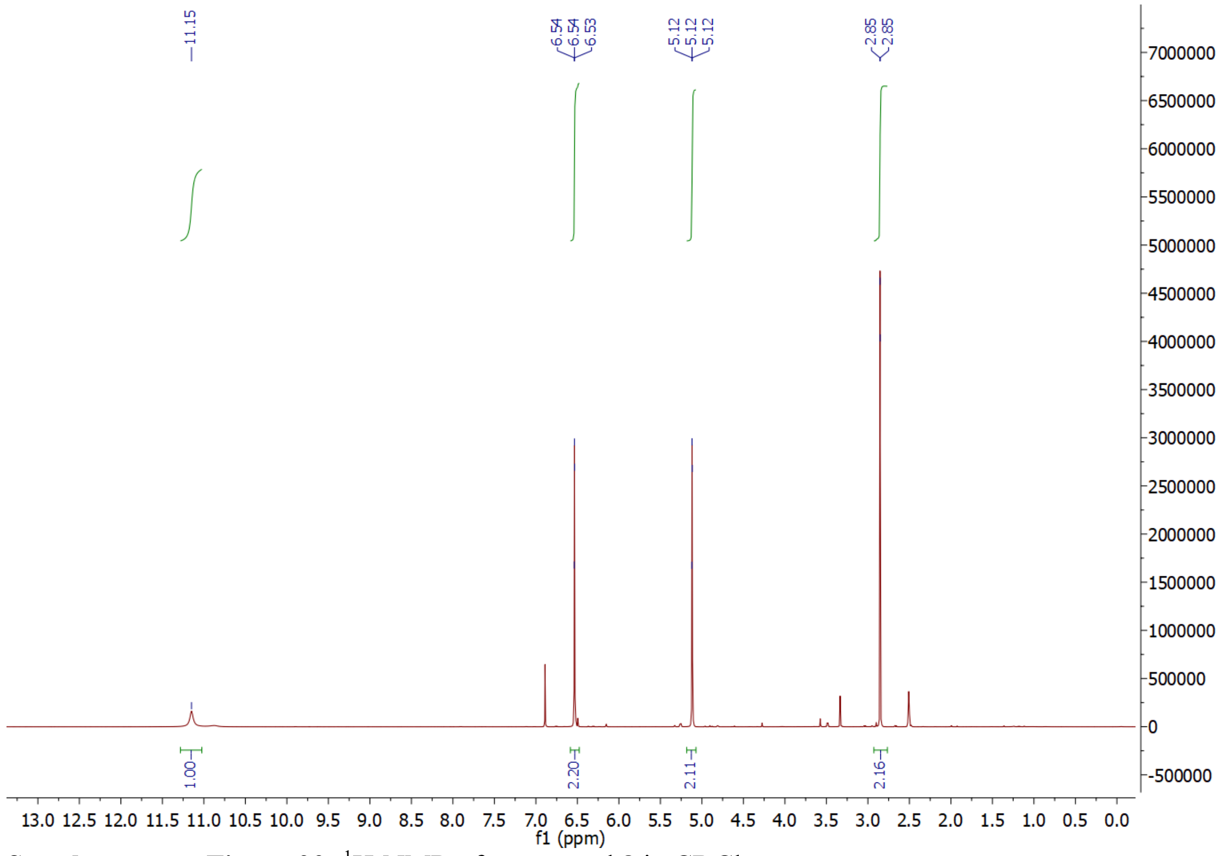


**Supplementary Figure 23:** $^{1}H$-NMR of compound **8** in $CDCl_3$.

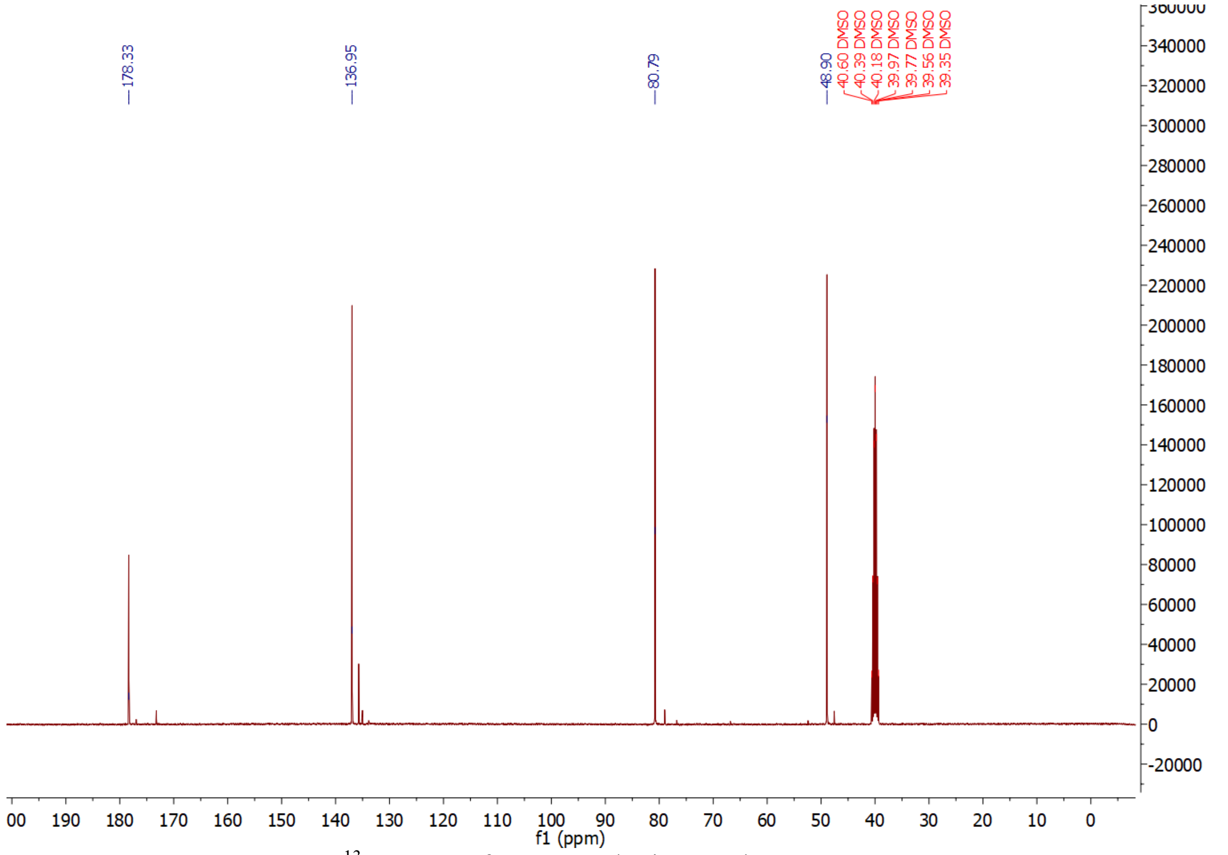


**Supplementary Figure 24:** $^{13}C$-NMR of compound **8** in $CDCl_3$.

**1-(4-((5-methoxy-2-nitro-4-(prop-2-yn-1-yloxy)benzyl)oxy)benzyl)-1H-pyrrole-2,5-dione (9):** **8** (850 mg, 5.1 mmol, 1.1 Eq.), 7 (1.94 g, 4.77 mmol, 1.0 Eq.) and potassium carbonate (3.37 g, 24.4 mmol, 5.1 Eq.) were added to DMF (50 mL) and the reaction mixture was stirred for 2 h at 90 °C. The mixture was than diluted with water (100 mL) and extracted with EtOAc (3 x 80 mL). The organic phase was dried with magnesium sulfate and the solvent was removed in vacuo. The residue was heated to 150 °C at 2 mbar for 1 h and afterwards the crude product was purified by column chromatography on silica (1:1 EtOAc/Pentane). The final compound (252 mg, 0.59 mmol, 13%) was yielded as light yellowish solid. $^{1}$H-NMR (300 MHz, DMSO): $\delta$ = 7.85 (s, 1H), 7.35 (s, 1H), 7.19 (d, $J$ = 8.6 Hz, 2H), 7.05 (s, 2H), 6.99 (d, $J$ = 8.6 Hz, 2H), 5.39 (s, 2H), 4.96 (d, $J$ = 2.4 Hz, 2H), 4.53 (s, 2H), 3.88 (s, 3H), 3.66, (t, $J$ = 2.4 Hz, 1H). $^{13}$C-NMR (75 MHz, DMSO): $\delta$ = 171.3, 157.8, 154.0, 145.9, 139.9, 135.1, 130.0, 129.4, 128.7, 115.4, 112.1, 110.9, 79.7, 78.9, 67.1, 57.0, 56.8. HR-MS (ESI): m/z calculated for $C_{22}H_{18}N_2O_7Na$ $[M+Na]^+$: 445.1003, found: 445.1006. IR (ATR) [cm$^{-1}$]: 2357, 2335, 1705, 1521, 1507, 1387, 1320, 1286, 1243, 1215, 1072, 821, 693. Mp.: 152–154 °C.

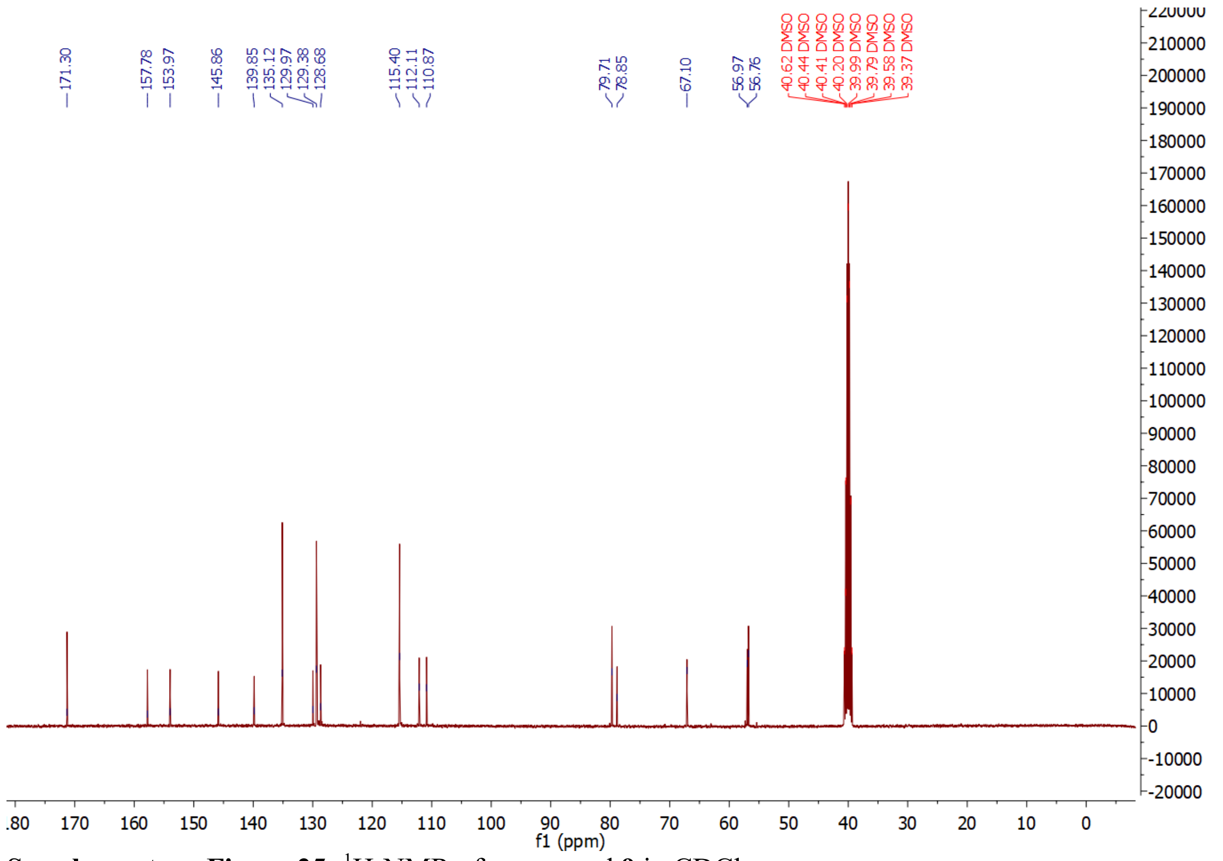


**Supplementary Figure 25:** $^{1}$H-NMR of compound **9** in $CDCl_3$.

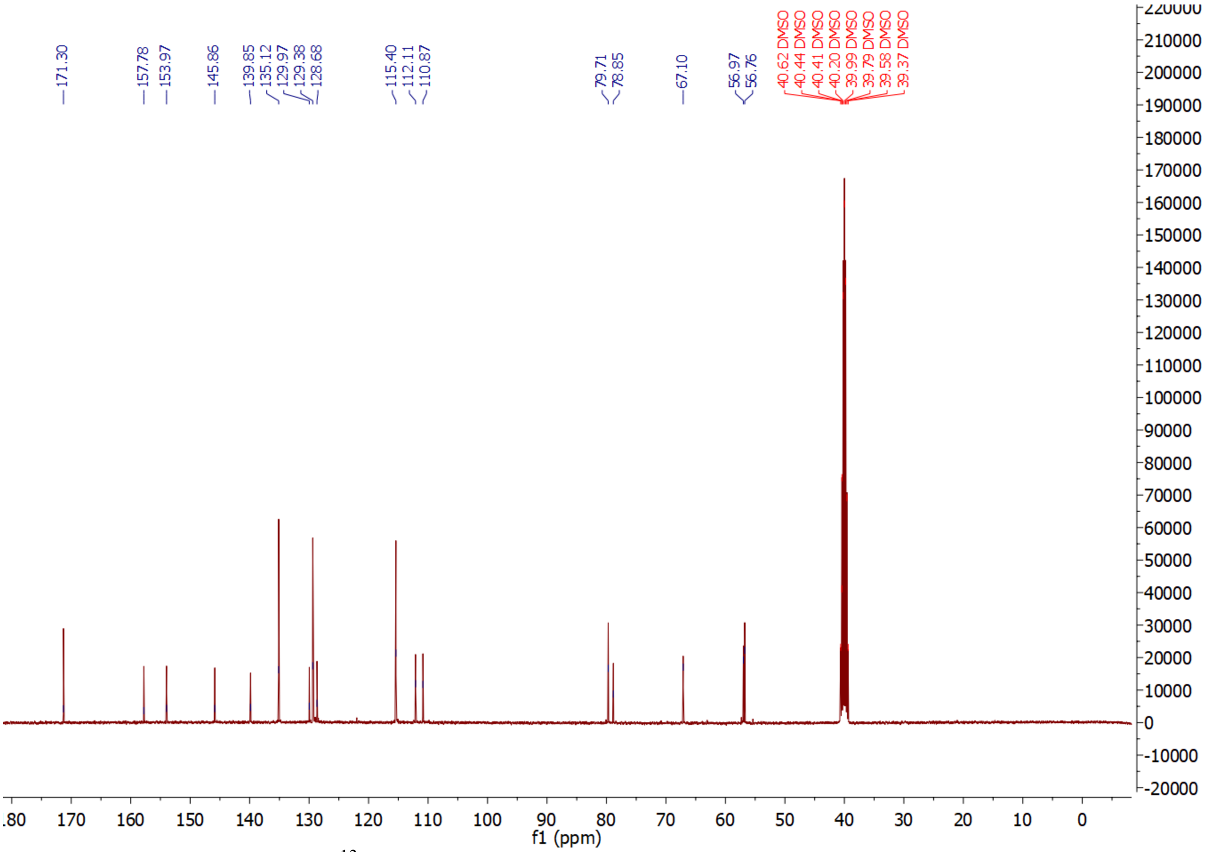


**Supplementary Figure 26:** $^{13}$C-NMR of compound **9** in $CDCl_3$.

**2-(4-((4-((4-((2,5-dioxo-2,5-dihydro-1H-pyrrol-1-yl)methyl)phenoxy)methyl)-2-methoxy-5-nitrophenoxy)methyl)-1H-1,2,3-triazol-1-yl)acetic acid (11): 9** (20 mg, 0.05 mmol, 1.0 Eq) was dissolved in DMSO (750 µL) in a crimp-top vial. azidoacetic acid (4 µL, 0.05 mmol, 1 Eq), $CuSO_4$ (0.25 M solution in MilliQ, 4 µL, 0.001 mmol, 0.02 Eq), Na-ascorbate (1.0 M solution in MilliQ, 5 µL, 0.005 mmol, 0.1 Eq) and MilliQ (250 µL) were added, and the reaction mixture was left stirring at room temperature for 3 hours.Precipitation of the desired product indicated completion of the reaction. The reaction mixture was then centrifuged, the supernatant was removed, and the precipitate was redissolved in MeCN. The solvent was then evaporated *in vacuo*. To eliminate any residual DMSO, the compound was lyophilized overnight. Compound **11** was obtained as a pale yellow solid (26.2 mg, quantitative yield) and converted without further purification. LCMS-ESI$^+$ (m/z): calcd. for $C_{24}H_{21}N_5O_9$ $[M+H]^+$ 524.13; found: 524.1.

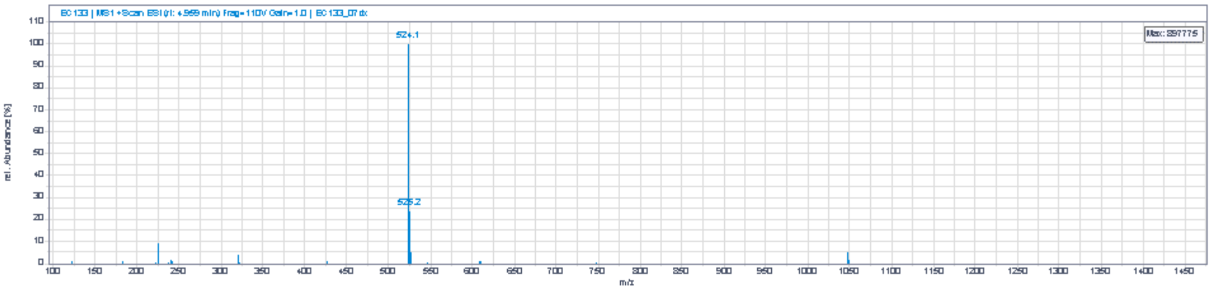


**Supplementary Figure 27:** LCMS of compound **11**.

**ALFA peptide:** The ALFA peptide used to synthesize the **LRT** was synthesized *via* microwave-assisted solid phase peptide synthesis on a Liberty Blue Peptide Synthesizer (CEM) following SOP1. The synthesis (on a 0.025 mmol scale) was carried out on a preloaded Fmoc-Glu-Wang resin (0.32 mmol/g loading). DIC (0.25 M in DMF) and Oxyma (0.50 M in DMF) were employed as activators, while deprotection was carried out with a 20% solution of piperidine in DMF.

**Supplementary Table 13**: Amino Acid building blocks and scale used for SPPS.

| Amino Acid | Molecular Weight (g/mol) | Concentration (M) | Mass (g) | DMF Volume (mL) |
|---|---|---|---|---|
| Fmoc-Arg(Pbf)-OH | 311.33 | 0.2 | 0.50 | 8 |
| Fmoc-Gly-OH | 353.41 | 0.2 | 0.43 | 6 |
| Fmoc-Glu(OtBu)-OH | 425.47 | 0.2 | 0.36 | 4 |
| Fmoc-Lys(Boc)-OH | 468.54 | 0.2 | 0.29 | 4 |
| Fmoc-Pro-OH | 337.37 | 0.2 | 0.14 | 2 |
| Fmoc-Ser(OtBu)-OH | 383.44 | 0.2 | 0.24 | 3 |
| Fmoc-Thr(tBu)-OH | 468.54 | 0.2 | 0.24 | 3 |

After the synthesis, the resin was collected in a BD-Syringe and cleavage was performed according to SOP2. A small aliquot was re-dissolved in MeOH for ESI-MS analysis. HR-MS (ESI$^+$) – m/z calculated for $[C_{96}H_{151}N_{29}O_{31}]$: 2206.1; found: m/z 2206.1 [M]; m/z 1104.5672 $[M+2H]^{2+}$; m/z 736.7135 $[M+3H]^{3+}$; m/z 552.7871 $[M+4H]^{4+}$.

**Light Responsive Tag (LRT):** The synthesis of the LRT is described at the beginning of the Supplementary Information.

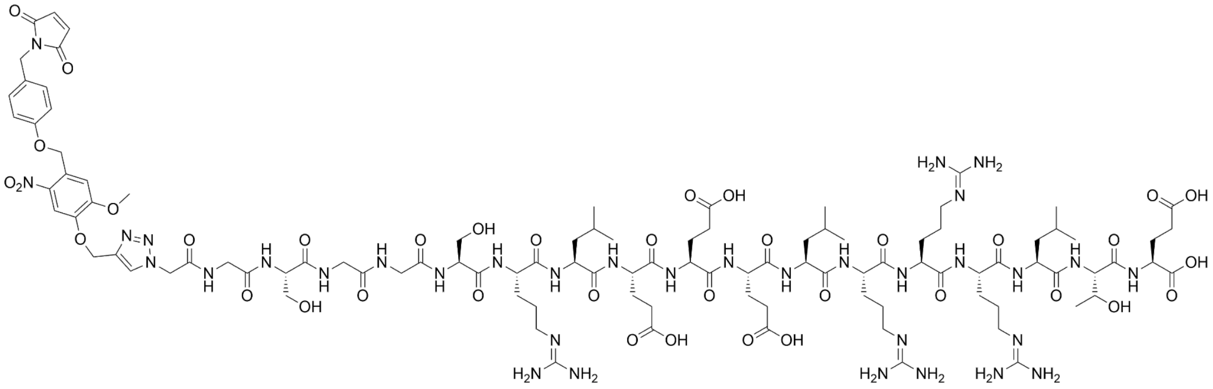

**Supplementary Figure 28:** Chemical structure of the LRT.

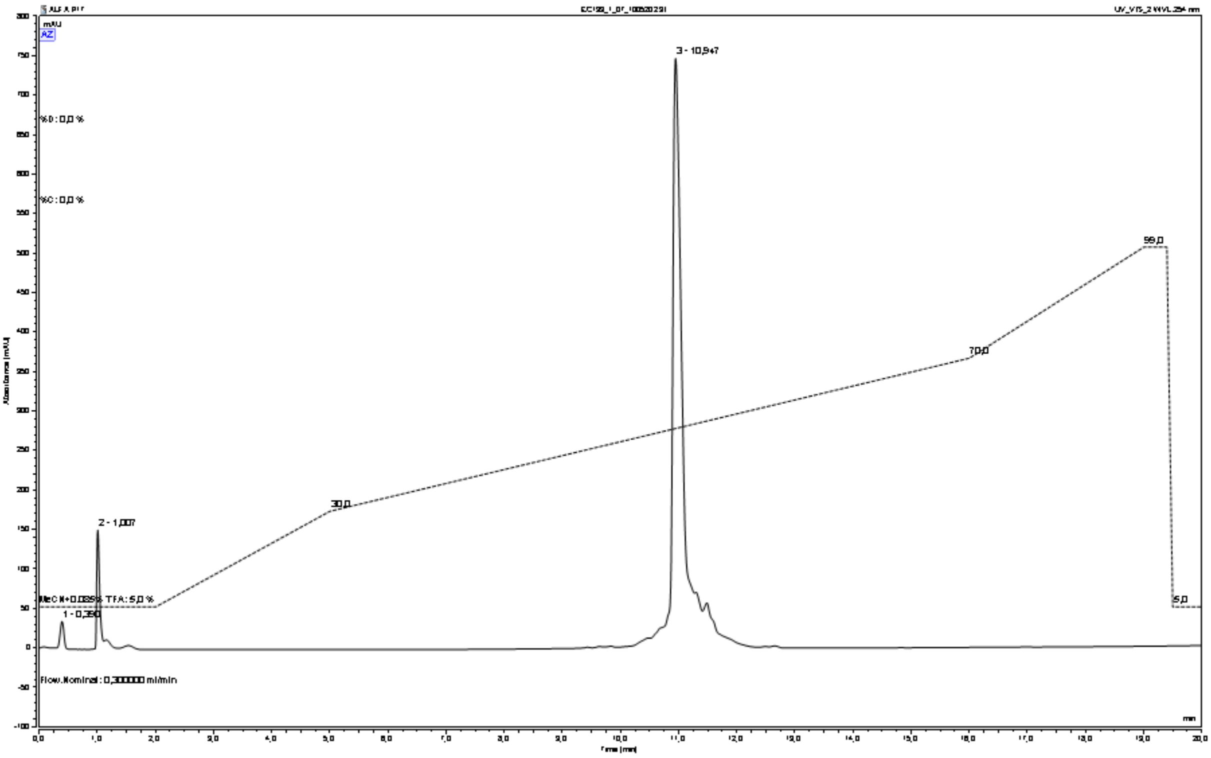


**Supplementary Figure 29:** UPLC-trace of LRT at 254 nm and solvent gradient used for elution.

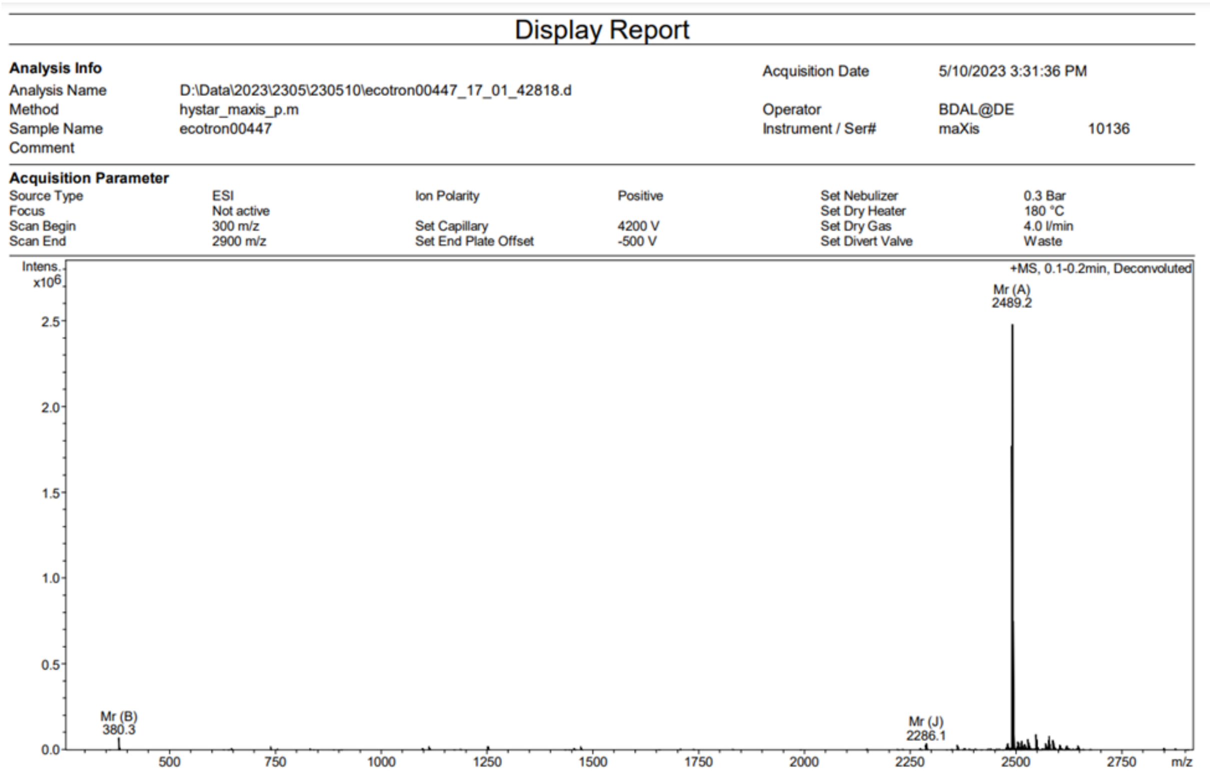

### Display Report

**Analysis Info**

| | | | |
|---|---|---|---|
| Analysis Name | D:\Data\2023\2305\230510\ecotron00447_17_01_42818.d | Acquisition Date | 5/10/2023 3:31:36 PM |
| Method | hystar_maxis_p.m | Operator | BDAL@DE |
| Sample Name | ecotron00447 | Instrument / Ser# | maXis 10136 |
| Comment | | | |

**Acquisition Parameter**

| | | | | | |
|---|---|---|---|---|---|
| Source Type | ESI | Ion Polarity | Positive | Set Nebulizer | 0.3 Bar |
| Focus | Not active | | | Set Dry Heater | 180 °C |
| Scan Begin | 300 m/z | Set Capillary | 4200 V | Set Dry Gas | 4.0 l/min |
| Scan End | 2900 m/z | Set End Plate Offset | -500 V | Set Divert Valve | Waste |

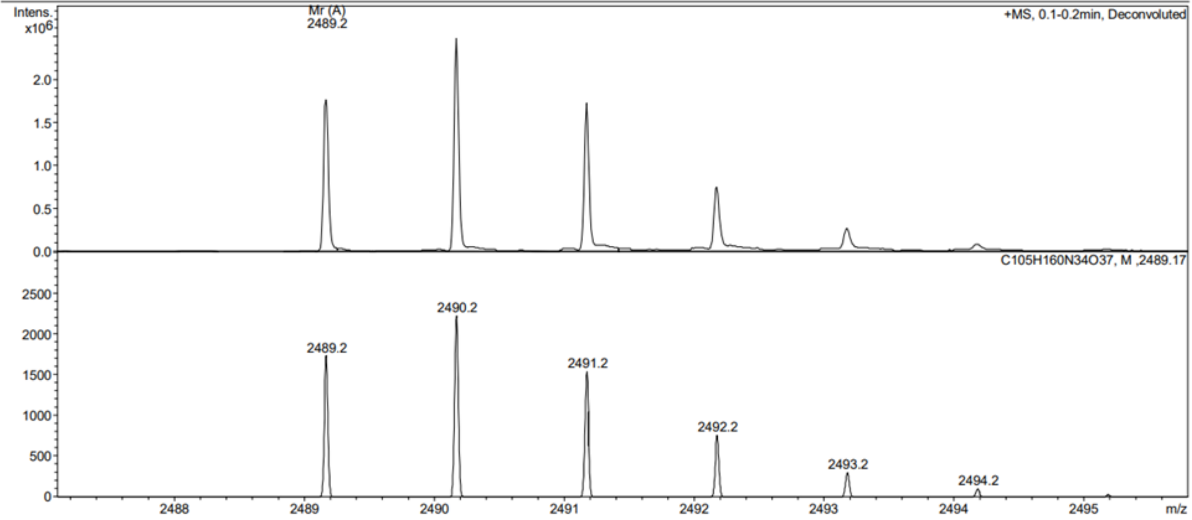


1

### Display Report

**Analysis Info**

| | | | |
|---|---|---|---|
| Analysis Name | D:\Data\2023\2305\230510\ecotron00447_17_01_42818.d | Acquisition Date | 5/10/2023 3:31:36 PM |
| Method | hystar_maxis_p.m | Operator | BDAL@DE |
| Sample Name | ecotron00447 | Instrument / Ser# | maXis 10136 |
| Comment | | | |

**Acquisition Parameter**

| | | | | | |
|---|---|---|---|---|---|
| Source Type | ESI | Ion Polarity | Positive | Set Nebulizer | 0.3 Bar |
| Focus | Not active | | | Set Dry Heater | 180 °C |
| Scan Begin | 300 m/z | Set Capillary | 4200 V | Set Dry Gas | 4.0 l/min |
| Scan End | 2900 m/z | Set End Plate Offset | -500 V | Set Divert Valve | Waste |

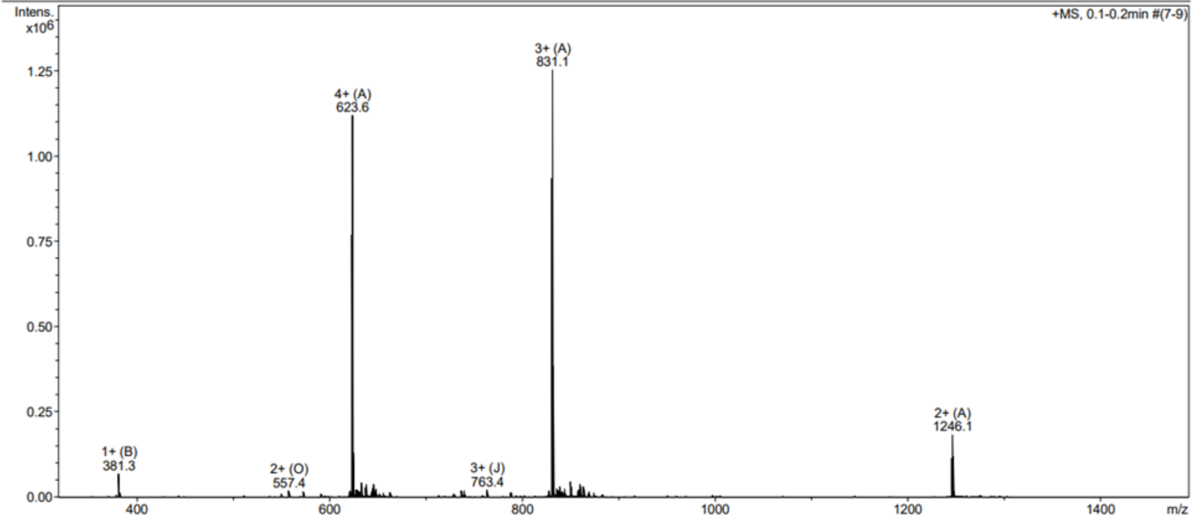


2

### Display Report

**Analysis Info**

| | | | |
|---|---|---|---|
| Analysis Name | D:\Data\2023\2305\230510\ecotron00447_17_01_42818.d | Acquisition Date | 5/10/2023 3:31:36 PM |
| Method | hystar_maxis_p.m | Operator | BDAL@DE |
| Sample Name | ecotron00447 | Instrument / Ser# | maXis 10136 |
| Comment | | | |

**Acquisition Parameter**

| | | | | | |
|---|---|---|---|---|---|
| Source Type | ESI | Ion Polarity | Positive | Set Nebulizer | 0.3 Bar |
| Focus | Not active | | | Set Dry Heater | 180 °C |
| Scan Begin | 300 m/z | Set Capillary | 4200 V | Set Dry Gas | 4.0 l/min |
| Scan End | 2900 m/z | Set End Plate Offset | -500 V | Set Divert Valve | Waste |

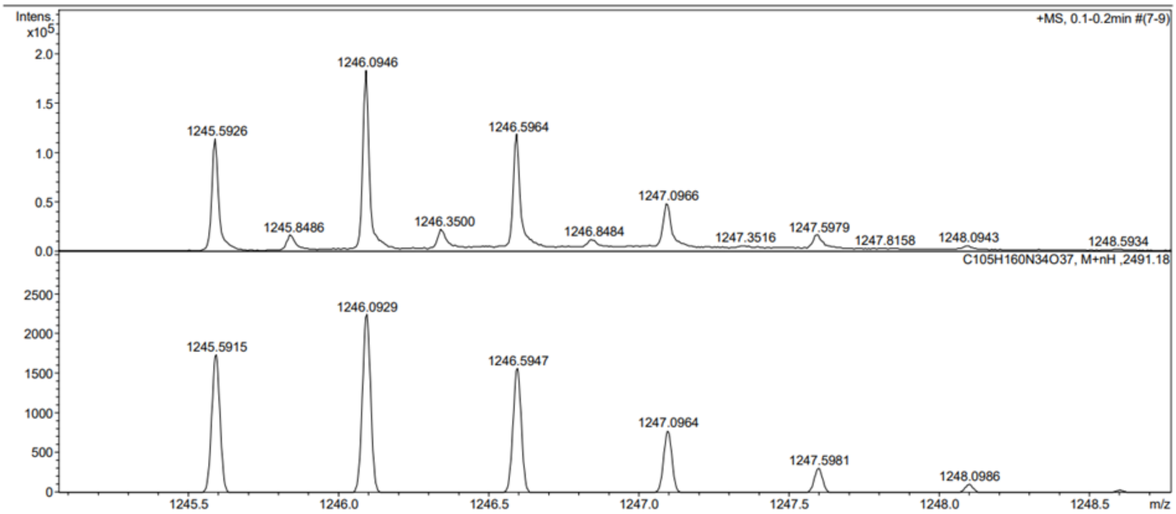


1

### Display Report

**Analysis Info**

| | | | |
|---|---|---|---|
| Analysis Name | D:\Data\2023\2305\230510\ecotron00447_17_01_42818.d | Acquisition Date | 5/10/2023 3:31:36 PM |
| Method | hystar_maxis_p.m | Operator | BDAL@DE |
| Sample Name | ecotron00447 | Instrument / Ser# | maXis 10136 |
| Comment | | | |

**Acquisition Parameter**

| | | | | | |
|---|---|---|---|---|---|
| Source Type | ESI | Ion Polarity | Positive | Set Nebulizer | 0.3 Bar |
| Focus | Not active | | | Set Dry Heater | 180 °C |
| Scan Begin | 300 m/z | Set Capillary | 4200 V | Set Dry Gas | 4.0 l/min |
| Scan End | 2900 m/z | Set End Plate Offset | -500 V | Set Divert Valve | Waste |

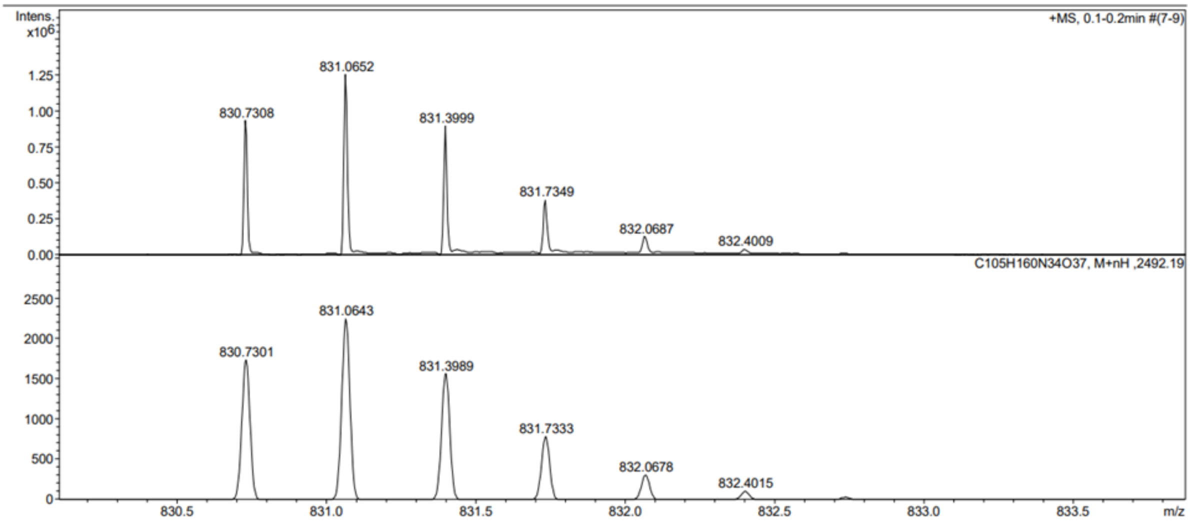


2

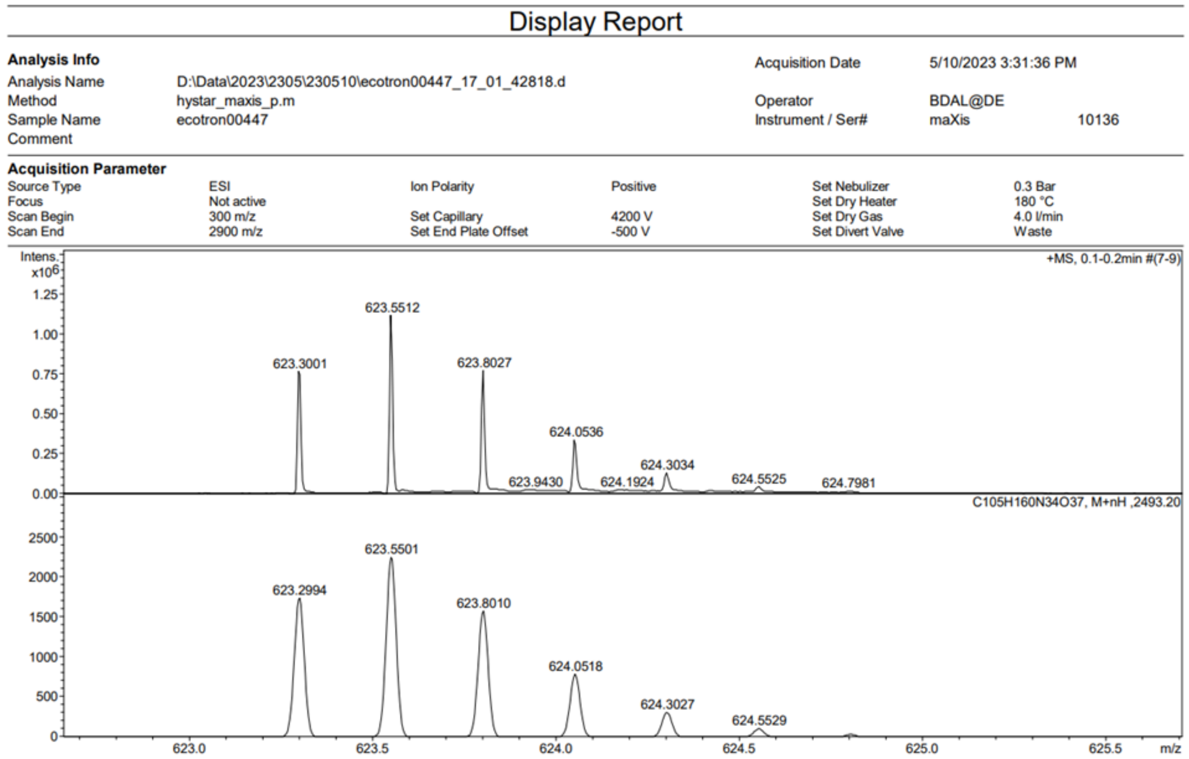


**Supplementary Figure 30:** High-Resolution Mass Spectroscopy (HR-MS) spectra of the LRT.

**References**

1. Stranius, K. & Börjesson, K. Determining the Photoisomerization Quantum Yield of Photoswitchable Molecules in Solution and in the Solid State. *Sci. Rep.* **7**, 41145 (2017).
